# Endothelial Immunosuppression in Atherosclerosis : Translational Control by Elavl1/HuR

**DOI:** 10.1101/2024.08.02.605922

**Authors:** Sarah-Anne E. Nicholas, Stephen R. Helming, Antoine Ménoret, Christopher Pathoulas, Maria M. Xu, Jessica Hensel, Amy L. Kimble, Brent Heineman, Evan R. Jellison, Bo Reese, Beiyan Zhou, Annabelle Rodriguez-Oquendo, Anthony T. Vella, Patrick A. Murphy

## Abstract

Atherosclerotic plaques are defined by the accumulation of lipids and immune cells beneath the endothelium of the arterial intima. CD8 T cells are among the most abundant immune cell types in plaque, and conditions linked to their activation correlate with increased levels of cardiovascular disease. As lethal effectors of the immune response, CD8 T cell activation is suppressed at multiple levels. These checkpoints are critical in dampening autoimmune responses, and limiting damage in cardiovascular disease.

Endothelial cells are well known for their role in recruiting CD8 T and other hematopoietic cells to low and disturbed flow (LDF) arterial regions that develop plaque, but whether they locally influence CD8 effector functions is unclear. Here, we show that endothelial cells can actively suppress CD8 T cell responses in settings of chronic plaque inflammation, but that this behavior is governed by expression of the RNA-binding protein Embryonic Lethal, Abnormal Vision-Like 1 (Elavl1). In response to immune cell recruitment in plaque, the endothelium dynamically shifts splicing of pre-mRNA and their translation to enhance expression of immune-regulatory proteins including C1q and CD27. This program is immuno-suppressive, and limited by Elavl1. We show this by *Cdh5(PAC)-CreERT2*-mediated deletion of Elavl1 (ECKO), and analysis of changes in translation by Translating Ribosome Affinity Purification (TRAP). In ECKO mice, the translational shift in chronic inflammation is enhanced, leading to increased ribosomal association of C1q components and other critical regulators of immune response and resulting in a ∼70% reduction in plaque CD8 T cells. CITE-seq analysis of the remaining plaque T cells shows that they exhibit lower levels of markers associated with T cell receptor (TCR) signaling, survival, and activation. To understand whether the immunosuppressive mechanism occurred through failed CD8 recruitment or local modulation of T cell responses, we used a novel *in vitro* co-culture system to show that ECKO endothelial cells suppress CD8 T cell expansion—even in the presence of wild-type myeloid antigen-presenting cells, antigen-specific CD8 T cells, and antigen. Despite the induction of C1q mRNA by T cell co-culture in both wild-type and ECKO endothelial cells, we find C1q protein abundantly expressed only in co-culture with ECKO cells. Together, our data define a novel immune-suppressive transition in the endothelium, reminiscent of the transition of T cells to T-regs, and demonstrate the regulation of this process by Elavl1.

## Introduction

Atherosclerotic lesion formation is initiated by endothelial cell (EC) activation and dysfunction in areas of arteries exposed to low and disturbed blood flow (LDF) – and not steady laminar flow^1–3^. In response to LDF, changes in aortic EC transcriptional programming facilitate immune cell recruitment and infiltration of the arterial intima^4,5^. The signals regulating this recruitment process have been well described, and transcriptional networks have been defined *in vitro*^6^ and more recently *in vivo*^7–9^. Major contributors to acute recruitment include adhesion molecules upregulated in ECs (e.g., P-selectin, ICAM1, and VCAM) and secreted cytokines (e.g., CCL2, CCL5) organized by transcriptional networks including elevated p65/NF-kB and reduced KLF2^10,11^, and also cytoskeletal and extracellular matrix changes mediated by integrin alpha5 and ‘response-to-injury’ matrix proteins such as fibronectin^12–14^. Recruited cells include large numbers of myeloid and lymphoid cells, potentially setting the stage for autoimmune-like damage to the vascular wall.

The endothelium is also well positioned to regulate chronic interactions between immune cells and the vessel wall, but very little is known about this process. Antigen-presenting CD11c cells sit just beneath ECs and above the basement membrane, where they send processes into the lumen of the vessel across the endothelium^15,16^. Despite this proximity, and the expression of a wide repertoire of immune-regulatory proteins^17^, how ECs respond in these conditions to suppress or enhance immune cell responses is uncharted territory.

Understanding whether ECs restrain unwanted immune cell activation is essential for understanding plaque progression, because immune cell infiltration is a common feature of the largest risk factors for cardiovascular disease, including hypercholesterolemia, aging, autoimmune disease, and viral infection.

Each of these conditions drives systemic activation of circulating T cells^18,19^, which are preferentially recruited to sites of LDF. CD8 T effector cells have a prominent role in the plaque, where they are retained after recruitment for months to years, poised for the secretion of cytotoxic and destructive proteins like granzyme K and perforin^20,21^. Previously, we demonstrated changes in splicing of pre-mRNA encoding proteins involved in mesenchymal-like and immune transitions of ECs under LDF, and identified a large set of regulatory RNA-binding proteins, including Elavl1/HuR and PTBP1^14,22–24^. These programs respond directly to the recruitment of myeloid and lymphoid cells to the arterial wall, suggesting that they could contribute to EC processes affecting local immune cell activation. However, with a few exceptions^14,23^, the extent to which these alternatively spliced pre-mRNA transcripts are translated into protein is still unclear. More importantly, it is unclear how this process affects immune cell physiology in the plaque.

Here, we use <u>T</u>ranslating <u>R</u>ibosome <u>A</u>ffinity <u>P</u>urification (TRAP) to examine the chronic adaptation of ECs in the atherosclerotic intima. We show that Elavl1 is increased in sites of LDF, which restrains a checkpoint program in ECs, suppressing ribosomal recruitment of mRNA that encodes immunosuppressive proteins, like C1q, and limiting CD8 T cell numbers in plaque. These findings confirm a previous observation that mRNA encoding C1q subunits are increased in ECs under chronically inflamed LDF conditions^25,26^ but shows that mRNA expression is not sufficient, as the levels of Elavl1 determines the translation potential of this mRNA. Using a novel *in vitro* co-culture assay of aortic ECs, primary monocyte-derived antigen-presenting cells, and CD8 T cells, we show that CD8 T cells induced mRNA expression of C1q in endothelial cells, but that this is not sufficient for protein expression. Deletion of Elavl1 is required for C1q protein production in ECs, and the immunosuppression of CD8 T cell responses to antigen stimulation.

Together, our data identify a novel stromal checkpoint in atherosclerotic lesions, which requires expression of the RNA-binding protein Elavl1 in the endothelium and a modification of the translatome as a critical step towards CD8 T cell accumulation in atherosclerotic lesions and plaque development.

## Results

### Ribosomal profiling of endothelial cells in atherosclerotic lesions identifies changes in splicing and translation of immune-regulatory pathways

To examine the transcriptional changes in endothelial cells (EC) within well-developed atherosclerotic plaque, we employed a model of low flow induced by <u>P</u>artial <u>C</u>arotid <u>A</u>rterial <u>L</u>igation (PCAL) coupled with hyperlipidemia mediated by adenoviral delivery of mutated proprotein convertase subtilisin/kexin type 9 (PCSK9) (AAV-mPCSK9) and high fat diet^27^. Surgical ligation induces low flow along the length of the left carotid artery (LCA) while the contralateral right carotid artery (RCA) continues to experience normal flow, and thus serves as a valuable animal-specific contralateral control (Figure 1A).

**Figure 1.**
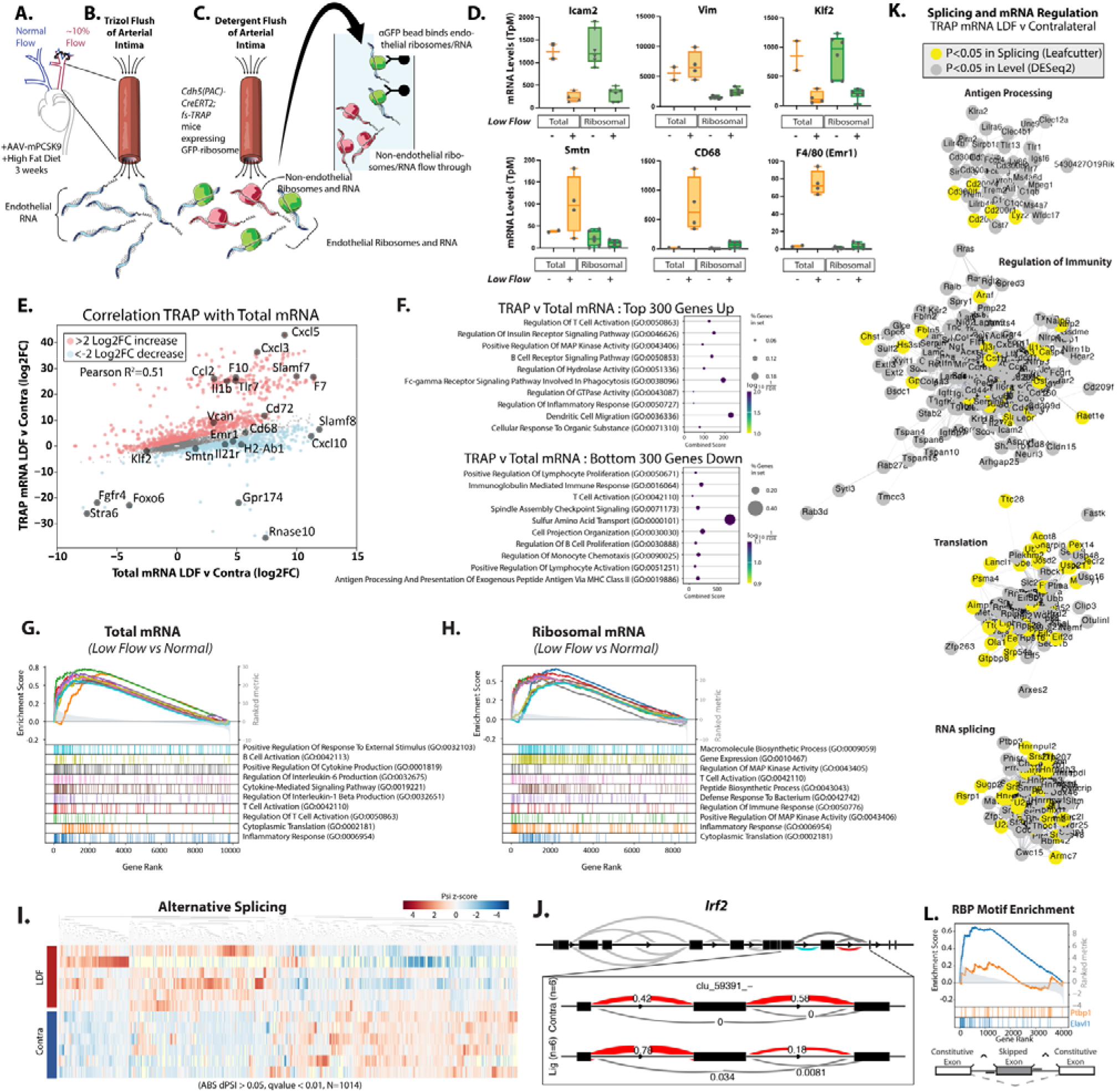
Translating ribosome affinity purification (TRAP) from atherogenic endothelium identifies changes in immune- regulatory pathways. (A) Partial Carotid Arterial Ligation (PCAL) induction of low flow in the left carotid artery (LCA) and hyperlipidemia collected at 3 weeks post PCAL. (B) Approach for isolating total intimal mRNA from both low and disturbed flow (LDF) and contralateral normal flow vessels via Trizol flush. (C) Approach for isolating endothelial-specific ribosomal mRNA from both low flow and normal flow vessels via RiboTRAP. (D) Expression level (FPKM) of indicated genes in low flow and normal flow vessels. (E) Plot showing the effects on transcript levels between LDF and contralateral arteries, between TRAP (Y-axis) and total mRNA (X-axis). (E) mRNA expression (Transcript per Million) detected in Total intimal RNAseq compared to TRAP RNAseq. (F-H) Gene Set Enrichment Analysis (GSEA) of low flow regulated transcripts thresholded on mean TpM greater than or equal to 100. Top 300 genes up indicate the genes and associated transcripts most enriched in TRAP low vs. contralateral relative to Total mRNA low vs. contralateral (red in E), down is depleted in TRAP vs. contralateral (blue in E). (G&H) GSEA plot of top 10 upregulated gene sets in GoTerms in (G) total mRNA and (H) ribosomal mRNA. (I&J) Leafcutter analysis of alternative splicing in ribosomal associated mRNA, showing clustering of groups by alternative splicing levels (I), where dPsi indicated the change in inclusion frequency between low flow and contralateral artery, plotted here as Z-scores and an example plotted splicing event (J). (K) Cytoscape visualization of top categories associated with low flow regulated transcripts in ribosomal mRNA. (L) Enrichment of Cis-RNAbp motifs nearby regulated skipped exons, showing the relative enrichment for Elavl1 and Ptbp1 6mers. (A-L) Total intimal RNAseq (Total; N=4 WT Ligated vs. N=2 Contralateral) compared to TRAP RNAseq (TRAP; N=6 WT Ligated vs. N=6 Contralateral) (K) Grey = differential mRNA TRAP, DESeq2 p.adjusted <0.05, yellow = differential mRNA splicing Leafcutter p.adjusted <0.05.

Previously, this model has been used to assess short-term EC responses to LDF by rapid intimal mRNA isolation using a Trizol flush method^7,22,24^ (Figure 1B). Recently, cell analysis of PCAL arteries identified an immune-like phenotype in the endothelium after two weeks of LDF^28^. Importantly, significant plaque development occurs by three weeks^27^, at which point there is substantial interaction between the immune system and ECs. Therefore, we chose to examine transcriptional alterations in arterial ECs three weeks after the induction of disturbed flow.

To assess mRNA translation, we used TRAP for the EC-specific isolation of ribosomes and associated RNA *in situ*^29^. Inducible VE-Cadherin-Cre activity (*Cdh5-CreERT2; RiboTRAP*) results in the selective expression of a GFP-tagged L10a ribosomal protein in ECs, which can be immunoprecipitated with associated RNA (Figure 1C). Typically, this is done from whole tissue homogenate, but we adapted this protocol based on our experience with rapid intimal Trizol flush to quickly isolate highly purified ribosomal complexes from the arterial intima. This flush substantially increases the specificity of the endothelial mRNA isolation above whole tissue homogenate by providing two levels of purification. We validated this approach by examining a hallmark EC transcript, Icam2 (Figure 1D). As expected, the transcript was detected at high levels (∼500 TpM) even in the non-L10a-GFP immunoprecipitated flow through, and only slightly increased (∼1200 TpM) in L10a-GFP bead isolate (SI Figure 1). These levels are comparable to what we have previously observed in Trizol flush, which is already >99% endothelial transcript^22^. Levels were lower in the ligated artery, coinciding with increased vimentin and fibronectin (Vim and Fn1), and indicating transcriptional alterations associated with the endothelial-to-mesenchymal-like transition (EndoMT) (Figure 1D and SI Figure 1, P<0.001). We expected the TRAP method to have the greatest impact on eliminating contaminating transcripts from other cells in the LDF intima, including myeloid cells and infiltrating mural cells, which would otherwise appear in a Trizol flush of all mRNA. Therefore, we examined expression of Smtn as a marker of mural cells and CD68 and Emr1(F4/80) as markers of myeloid cells and compared the relative expression of these to levels we had observed in Trizol flush. All were depleted in the L10a-GFP pulldowns, relative to a Trizol flush of all intimal mRNA, based on transcripts per million (TpM) (Figure 1D, P<0.0001). Thus, TRAP effectively enriched EC transcripts.

In general, as indicated by expression of KLF2 (Figure 1D) and correlational analysis (Figure 1E, R^2^=0.51), expression changes in the ribosome-associated mRNA resembled those in total mRNA from Trizol flush. However, we noted that the responses did not tightly correlate, and that there were many transcripts relatively unchanged in total mRNA, but strongly changed in their ribosomal enrichment (e.g., Cxcl5, Cxcl3, F10, Il1b) (SI Table 1, Ribo_v_Total_Diff). Pathway analysis of the 300 transcripts preferentially enriched in TRAP ribosome-associated mRNA vs. Total mRNA indicates regulation of immune responses in both groups (Figure 1F). An examination of the overall transcriptional responses in the low flow atherosclerotic endothelium in Total and TRAP mRNA indicated similar pathways, notably also containing T cell activation (GO:0042110) (Figure 1G&H). We then analyzed alternative splicing changes in ribosome-bound RNA and identified a total of 3011 splicing changes (P<0.05) between LDF and normal flow contralateral arteries (Figure 1I, SI Table 2 Splice_Ribo_Lig_v_Contra). As an example, Irf2 exhibits a ∼40% increase in the inclusion of a skipped exon (Figure 1J). Irf2 is a critical regulator of MHC-I activity, and CD8 interactions and immune evasion in parenchymal cells^30^. Differential expression and splicing analysis of all significantly regulated LDF transcripts (P<0.05) showed an enrichment of genes associated with immunity, translation, RNA splicing, and antigen processing (Figure 1K and SI Figure 2).

**Figure 2.**
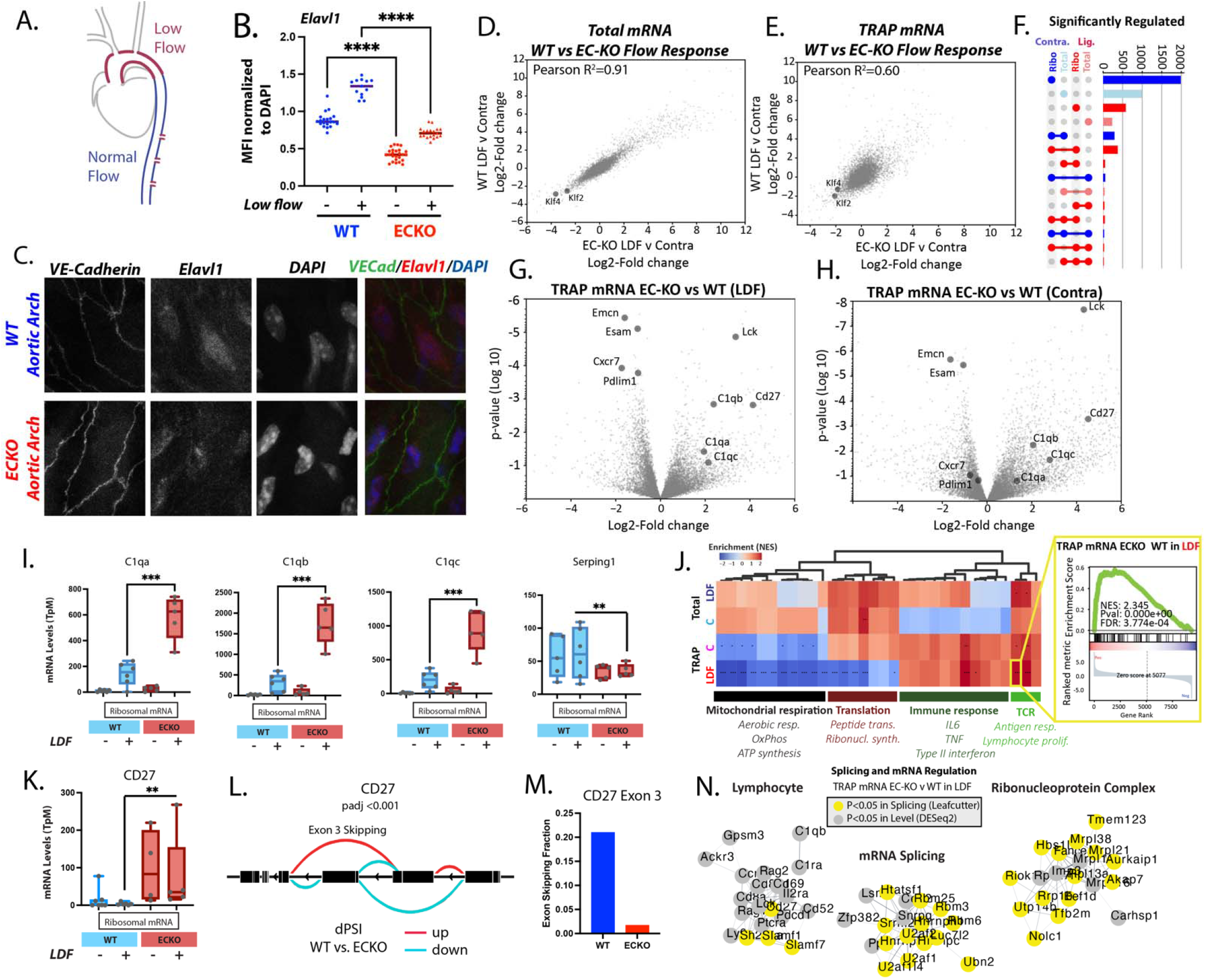
Endothelial Elavl1 is increased in atherogenic endothelium and regulates the translatome. (A) Schematic of regions of the aortic arch exposed to low and disturbed while the descending aorta is exposed to normal flow. (B&C) Elavl1 immunostaining in nuclei of aortic arch endothelial cells prepared en face. (B) Elavl1 mean fluorescence intensity (MFI) normalized to DAPI MFI for each nuclei. Each dot represents one nuclei from at least 3 separate fields of view. (C) Representative image, showing VE-cadherin in green, Elavl1 in red, DAPI in blue. (D&E) Plots showing correlations in gene expression between wild-type and ECKO carotid intima in the PCAL model, when total Trizol flushed mRNA is assessed (D) or TRAP-mRNA is assessed (E). (F) Upset plot showing the overlap in gene expression (DESeq2 pvalue<0.05) changes between wild-type and ECKO mice, in the low flow (LDF) artery, or the contralateral artery (C), when assessing either total mRNA (Total) or TRAP mRNA (Ribo). (G&H) Volcano plot showing differences in gene expression in TRAP mRNA between wild-type and ECKO mice in the low flow artery (LDF) or the contralateral artery (Contra). (I) Example plots showing the mean expression level for select genes in the LDF and contralateral arteries of wild- type and ECKO mice. (J) Gene set enrichment analysis of pathways affected in total mRNA and TRAP mRNA between wild-type and ECKO mice, showing significantly enriched GO terms, clustered into major categories. Inset shows an example of the underlying data for each square on the heatmap. (K) Example expression change, and (L) Leafcutter plot for a transcript differentially spliced and expressed in TRAP mRNA. (M) Shows the absolute change in Exon 3 inclusive transcript. (N) Clustering of transcripts by Cytoscape, which are altered at expression level or splicing in the LDF endothelial cells, when comparing wild-type and ECKO mice. (B) ***p<0.0001, ANOVA with post-hoc Tukey test. (I&K) **p<0.001, ***p<0.0001, DEseq2. (J) *False Discovery Rate (FDR)<0.05, **FDR<0.01, ***FDR<0.001 by GSEA weighted analysis.

As splicing and translation are regulated by RNA-binding proteins, we sought to identify RNA-binding proteins (RBPs) involved in the LDF-responsive translatome and spliceosome. Therefore, we performed enrichment analysis around LDF-regulated alternatively spliced skipped exons in the TRAP isolate. We examined 6-mer RBP motifs enriched in the 200bp upstream or downstream of LDF-regulated skipped exons and compared these to skipped exons detected at a similar level but not regulated by LDF. The motif for RBP Elavl1, which we had previously identified as the motif most enriched in the acute (48hrs LDF) response ^22,24^, was among the top 5 motifs nearby splicing events changed in this chronic atherogenic state (3 wks LDF) (Figure 1L, P<0.0001). Notably, the Ptbp1 motif, which we had found was highly enriched nearby acutely changed splicing events (48 hrs LDF), was not significantly enriched here (3 wks LDF), suggesting differences in the regulation of acute and chronic responses (Figure 1L).

Thus, TRAP data identifies ribosome enrichment of transcripts predicted to affect immune-endothelial interactions, and motif analysis pinpoints Elavl1 as a key regulator of this response.

### The RNA-binding protein Elavl1 determines the splicing and ribosomal association of immune-regulatory pathways in plaque endothelium

To examine Elavl1 protein levels in the endothelium in sites of LDF, we performed immunostaining of LDF regions of the aortic arch and laminar flow regions of the descending aorta (Figure 2A). We found that Elavl1 protein was increased in ECs of the arch when compared to non-LDF sites in the descending aorta (Figure 2B&C). We previously observed that *Ptbp1* transcript increased in a platelet dependent manner in the acute response to disturbed flow. Similarly, *Elavl1* transcript induction after LDF required the presence of platelets (SI Figure 3). Analysis of public single-cell data showed that *Elavl1* mRNA is elevated in a subset of the endothelium under chronic LDF, two weeks after PCAL (SI Figure 4)^28^. Thus, Elavl1 may be an important regulator of the chronic adaptation of the endothelium to atherogenic conditions, and may regulate EC interactions with recruited immune cells.

**Figure 3.**
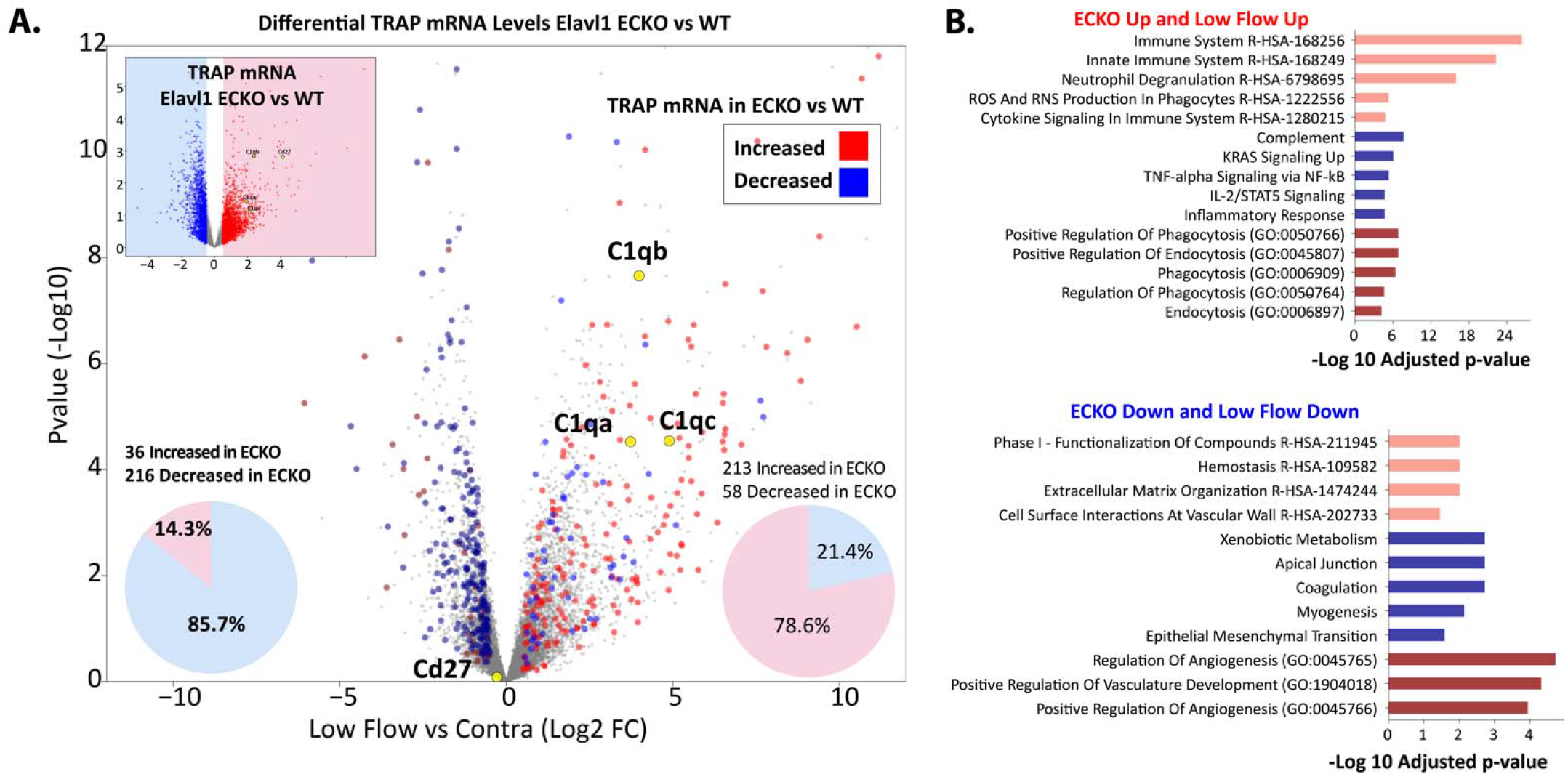
Deletion of endothelial Elavl1 enhances a subset of the endothelial translatome involved in immune regulation. (A, inset) Differential gene expression (DESeq2, Log2 Fold Change) in response to low flow, comparing Elavl ECKO to Elavl WT. Red represents transcripts enriched >0.5 Log2 Fold Change in ECKO TRAP mRNA vs. WT TRAP mRNA. Blue represents transcripts depleted <0.5 Log2 Fold Change in the same comparison. (A) Genes from (red and blue from A, inset) were plotted on a volcano plot showing differential TRAP mRNA expression the low flow artery and the contralateral artery in wild-type mice. Pie charts show the proportion of transcripts down (left) or up (right) in TRAP mRNA in low flow response that are down (blue) or up (red) in ECKO response. (B) Enrichr analysis of top gene categories where the natural low flow response is enhanced by the deletion of Elavl1 (216 genes decreased in the low flow TRAP mRNA and further decreased by the loss of Elavl1, or 213 genes increased in low from TRAP mRNA and further increased by loss of Elavl1). Pink = Reactome, Blue = msigDB Hallmark, Crimson = GO Terms.

**Figure 4.**
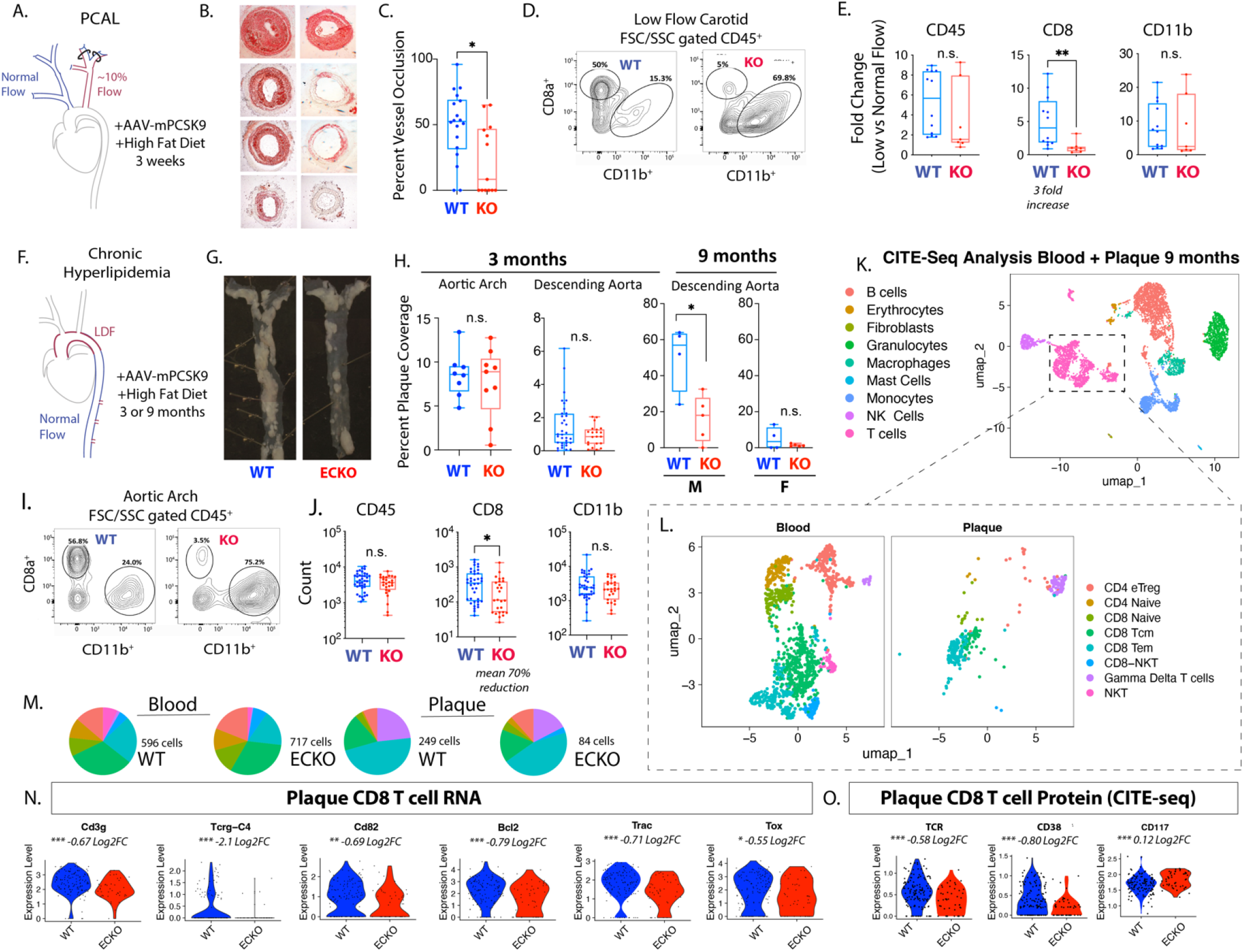
Endothelial deletion of Elavl1 reduces CD8 T cell numbers and activation in plaque progression. (A-C) Analysis of atherosclerotic plaque in ligated vessels 3 weeks post-PCAL; N=18 WT, N=12 EC-KO. (A) Schematic of PCAL model. (B) Examples of OilRedO stained cross-sections of the low flow LCA. (C) Quantification of plaque as percent vessel occlusion. (D) Flow Cytometry plots of size-gated CD45 cells isolated from atherogenic LCA. (E) Fold-increase in the low flow artery vs. contralateral normal flow control in the number of CD45+, CD45+CD8+, and CD45+CD11b+ cells. N=12 WT, N=7 EC-KO. (F-H) Analysis of atherosclerotic plaque in aortas of mice collected after 3 months or 9 months of hyperlipidemia. (G) Representative images of male aortas at 9 months hyperlipidemia. (H) Quantification of plaque as percent vessel coverage in the aortic arch and descending aorta. Males and females are separated in 9 month data, since differences could be observed at this time point in the hypercholesteremia induce (I&J) Flow Cytometry plots and quantitation of aortic arch after 3 months of hyperlipidemia. N=38 WT, N=26 EC-KO. (K-O) CITE-seq analysis of blood and aortic CD45+ cells at 9 months hyperlipidemia. ECKO and WT mice, N= 7 WT, N=7 ECKO. (K) Leiden clustering of all cells with annotation by RNA and protein labels, each dot represents one cell. (L) T cell subset, organized by tissue of origin. (M) Pie charts showing the total number and ratios of the T cell subsets in blood and in aortic tissue, colors as indicated in L. (N-O) Violin plots of gene expression (N, RNA) or protein expression (O, Protein CITE-seq) of plaque CD8 subcluster from (L). (C,E,H,J) *p<0.05, **p<0.01 by Mann-Whitney test. (N) Unadjusted p-values, *p<0.05, **p<0.01, p<0.001 by Student’s t-test, Log2FC from Seurat analysis.

To genetically define the role of EC Elavl1, we generated a novel, temporally inducible, EC-specific Elavl1 knockout (ECKO) mouse model (*Cdh5-CreERT2; Elavl1^f/f^*). Cre recombinase expression driven by VE-Cadherin (Cdh5) results in EC-specific excision of a region of Elavl1 containing the translation start site. Tamoxifen-induced recombination of mice at least 6 weeks old resulted in reduction of Elavl1 protein staining *ex vivo* (Figure 2B&C) and reduced levels of the excised region in transcripts from ECKO intimal RNA (*Cdh5-CreERT2; Elavl1^f/f^)* relative to WT littermate controls (*Elavl1^f/f^*) (SI Figure 5). We isolated mRNA from ECKO mice in the PCAL+AAV-PCSK9 model with either total mRNA flush or TRAP isolation. Overall, total mRNA expression changes of atherogenic endothelium were comparable in ECKO mice and controls (Figure 2D, Pearson correlation 0.91). Although Elavl1 can regulate mRNA stability^31^, loss of Elavl1 did not substantially affect total mRNA levels or the levels of canonically stabilized transcripts in this model (Figure 2D, SI Figure 6). However, major differences occurred at the level of ribosome association, where the response of Elavl1 ECKO mice diverged from wild-type mice (Figure 2E, Pearson correlation 0.6). For example, in ribosome-bound mRNA, we observed differences in transcripts from 138 genes (padj<0.05, N=5 ECKO v 6 WT) while observing only one differentially expressed transcript in total mRNA (N=4 ECKO v 4 WT). Relaxing significance thresholds to unadjusted p-value of 0.01, or 0.05 held the same trends (Figure 2E, 2F, SI Figure 7 and SI Table 1). Splicing, by Leafcutter analysis, similarly showed more transcripts affected by the deletion of Elavl1 in TRAP-mRNA (355 transcripts with padj<0.05) than in total mRNA (18 transcripts with padj<0.05) (SI Figure 8 and SI Table 2). Most genes were affected by either splicing or transcriptional changes, and typically not both (138 transcripts regulated at both splicing and expression levels of the 3011 regulated by splicing and 1019 by expression) (SI Figure 8). Thus, the major impact of Elavl1 is on the ribosome-associated mRNA.

**Figure 5.**
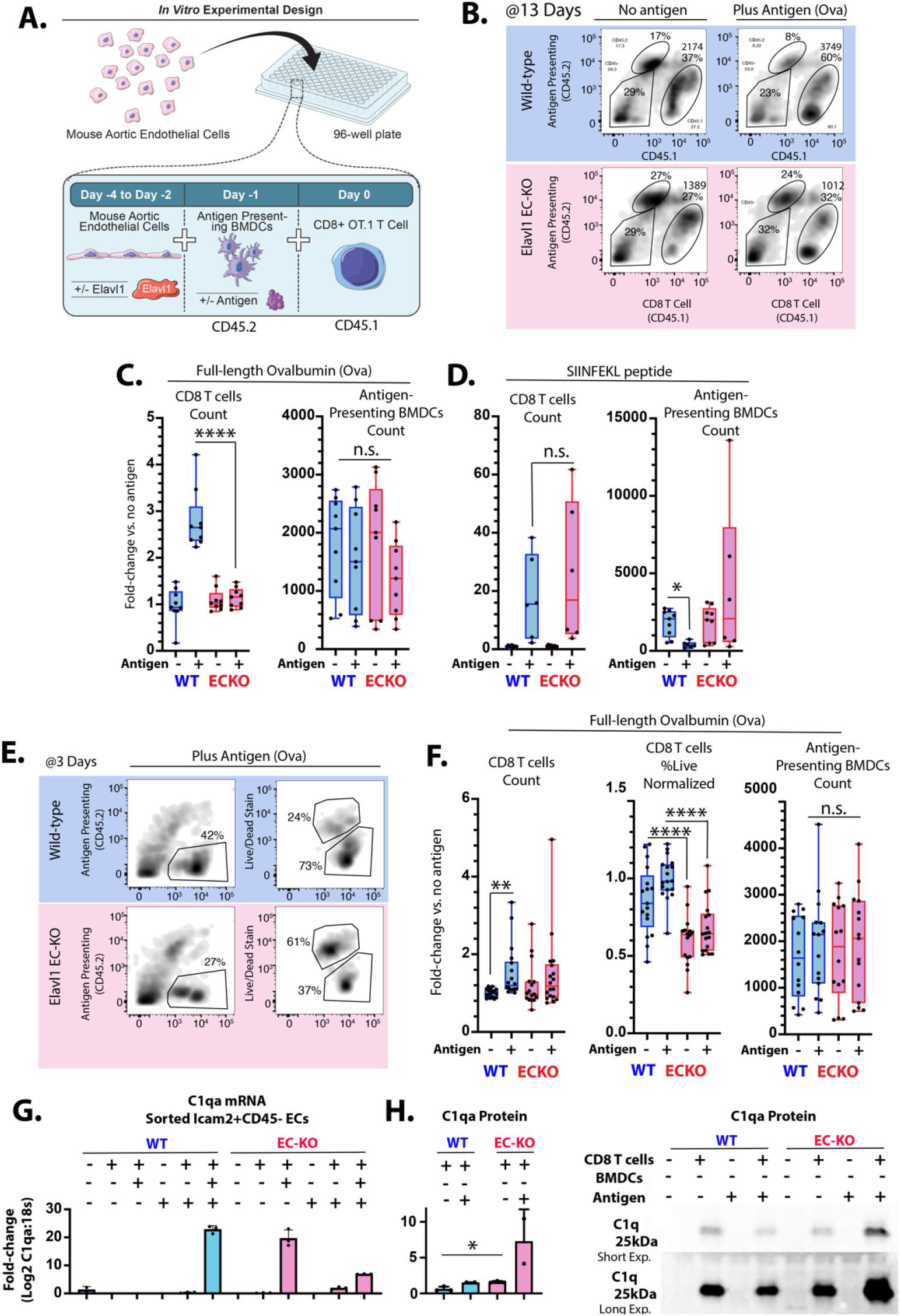
Deletion of Elavl1 from endothelial cells reduces short-term survival and long-term persistence of antigen responses in vitro. (A) Schematic of EC & BMDCs & CD45.1 OT.1 CD8 T cell co-culture in vitro. (B) Example flow cytometry plots of CD45.1 OT.1 T cells, after 13 days of co-culture. (C&D) Quantitation of CD45.1 CD8 OT.1 T cells and CD45.2 BMDCs after 13 days co- culture with or without cognate antigens (C) ovalbumin (0.5μg/mL OVA, in media throughout co-culture) or (D) SIINFEKL (1μg/mL, pre-incubation with endothelial cells and BMDCs). (E) Example flow cytometry plots after 3 days of coculture, showing live/dead gating of CD8 T cells. (F) Quantitation of CD45.1 CD8 OT.1 T cells, percent live CD8 OT.1 T cells and CD45.2 BMDCs. (G) Graph showing quantitative PCR-measured levels of C1qa, relative to 18s in endothelial cells sorted after 19 days of co-culture by Icam2+ and CD45neg from co-cultures with the indicated cell types. (H) Western blot showing the presence of C1q staining in equal amounts of media from the indicated co-culture conditions, with short and long exposure. (C,D,F,H) Each point represents results from an individual well. (C,D) N=9 vs 9, two separate experimental dates, (F) N>16 each, five experimental dates. (C,D,F) Data normalized to no antigen, wild-type ECs (except raw count data for BMDCs, which is not normalized). Comparisons by ANOVA with Sidaks multiple comparisons test, padj <0.05 = *; padj <0.005 = **, padj <0.0005 = ***, padj <0.0005 = ****.

Analysis of Elavl1 ECKO TRAP-mRNA, in both the LDF artery and the contralateral artery, revealed similar enrichment of transcripts associated with immunity (Figure 2G&H). Examples of increased transcript levels included *C1qa, C1qb, C1qc, CD27*, and *Lck* (Figure 2G–I), while decreased transcripts included *Emcn, Esam, Cxcr7*, and *Pdlim1* (Figure 2G&H). Pathway analysis indicated a suppression of transcripts associated with mitochondrial respiration and translation (Figure 2J and SI Figure 8), and an enrichment of genes modulating the immune response and TCR signaling (Figure 2J). An example <u>G</u>ene <u>S</u>et <u>E</u>nrichment <u>A</u>nalysis (GSEA) plot shows enriched terms for T cell activation pathways (inset from Figure 2J). Although most transcripts were affected by either expression or splicing, an exception was CD27 (SI Figure 9), an important co-stimulatory molecule, which showed increased levels of expression (Figure 2K) and increased splicing leading to exon 3 inclusive transcript in ECKO TRAP mRNA (Figure 2 L&M). This exon inclusion is predicted to extend the extracellular region of the protein to produce the full-length version of the protein, and the splicing change likely further increases the level of active CD27. A possibly similar switch between full length CD27 protein and a shorter variant has been observed in activated T cells^32^. The function of this altered splicing remains unclear, but human mutations affecting the alternatively spliced exon 3 are linked to immune dysfunction, suggesting possible activity in the regulation of immune responses^33^. Combining splicing and transcriptionally regulated gene sets showed common pathways affected by both processes (Figure 2N and SI Figure 10), again highlighting prominent effects on lymphocyte interactions and on mitochondrial respiration.

The Elavl1 ribosome-regulated T cell signature indicated enrichment of several transcripts typically thought of as lymphocyte-specific, such as Lck, Sla2, CD8a, CD8b, CD4, and Ptprc/CD45 (Figure 2N). Of these, Ptprc/CD45 is upregulated in ECs in EndoMT, where it contributes to the mesenchymal transition^34,35^. Lck, although well known for its functions in lymphocytes, is also expressed in the endothelium^36^. To our knowledge, expression of Sla2, CD8a, CD8b, and CD4 in the endothelium is unknown. In theory, this could represent contamination by non-endothelial transcript. In several instances, T cells have been shown to transfer material to other cells, including the epithelium and dendritic cells through exosomes^37,38^.

Regardless of the source, increased levels of transcript in the ribosomal fraction vs. the unbound flow through, indicates specific binding to the GFP-tagged EC ribosome (SI Figure 1 and SI Table 3, compare UnboundRibosomal to Ribosomal). Notably, these markers were also increased in the contralateral artery of Elavl1 ECKO mice where little immune cell recruitment is expected (Figure 2G and SI Tables 1&3).

To place the effect of Elavl1 deletion in the context of the change in ribosomal association which occurs in the LDF arterial endothelium, we examined the transcripts differentially increased (Figure 3A inset, red) or decreased (Figure 3A inset, blue) by Elavl1 to determine whether Elavl1 deletion suppressed or enhanced their ribosomal association in wild-type mice. We found that the vast majority (79%) of transcripts increased in ribosomal association under LDF were further increased by the loss of Elavl1 (Figure3A, red). Conversely, 86% of the transcripts less associated with the ribosome were further reduced by loss of Elavl1 (Figure 3A, blue). Thus, Elavl1 appears to limit a transition in mRNA translation that occurs in the atherogenic endothelium. Pathways associated with transcripts further increased by the loss of Elavl1 included the Immune System, Complement and Phagocytosis, while transcripts reduced included Hemostasis, Apical Junction and Angiogenesis (Figure 3B). C1q component transcripts C1qa, C1qb, and C1qc exemplifies this response. C1q expression increased only in the LDF endothelium and not under steady laminary flow in the contralateral artery (Figure 2J). Expression was further increased in ECKO TRAP mRNA (Figure 2J). The increase in endothelial C1q under chronic LDF resembles a similar change recently observed by single-cell RNA-seq^28^, and shows that Elavl1 regulates the ribosomal association of C1q mRNA, and the mRNA of other transcripts associated with the regulation of complement and immune responses (e.g., CD209d, Il2ra, IL17rb, Slamf1, CD27, and CD69), which differs from the effect on acute recruitment molecules (e.g. Icam1, Vcam, P- or E-selectin, Ccl2, or Ccl5). Together, our data implicates Elavl1 in the translational regulation of a specific subset of EC transcripts associated with immune regulation and not recruitment.

### Endothelium-specific deletion of Elavl1 reduced CD8 T cells and activation in plaque

To specifically examine the effects of EC Elavl1 deletion on endothelial interactions with immune cells in plaque, we tested two mouse models of atherosclerosis (Figure 4A&F). Both models developed plaque at sites of LDF, specifically in the ligated LCA 3 weeks after PCAL and hypercholesterolemia (Figure 4A–E) or at natural sites of LDF in the aorta after 3–9 months of chronic hypercholesterolemia (Figure 4F–O). We observed decreased levels of plaque in both models, with a mean carotid occlusion reduced from ∼50% in WT to ∼10% in ECKO mice in the PCAL model (Figure 4A–C, P<0.05), and reduced vessel plaque coverage in descending aortas of male mice from ∼50% in WT to ∼15% in ECKO after nine months of chronic hypercholesterolemia (Figure 4F–H, P<0.05). This occurred despite similarly increased cholesterol levels in WT and ECKO mice (SI Figure 11), although AAV-PCSK9 induced hypercholesterolemia and plaque development was generally reduced at late timepoints, as previously observed (SI Figure 10,11).

Notably, deletion of Elavl1 affected lesion progression, but not initial development. At 3 months, WT and ECKO mice had similar levels of plaque in the aortic arch and descending aorta, where only a few loci of plaque had begun to develop (Figure 4H). The primary difference at 3 months was the development of a few outliers in the top quartile of the male cohort with more extensive plaque (Figure 4H and SI Figure 12, fisher exact test P<0.01). Thus, although both WT and ECKO mice initially developed similar levels of plaque, these plaques grew faster in WT mice.

Given the prominent effects of Elavl1 deletion on transcriptional responses related to the immune response and T cell receptor signaling, we reasoned that T cell levels in atherosclerotic lesions might be impacted by Elavl1 expression. Murine ECs constitutively express MHC class I but not MHC class II molecules, allowing direct interactions with CD8 but not CD4 T cells^39,40^. Furthermore, CD8 T cells consistently associate with plaque development, so we chose to focus on them^20,21^. In the PCAL model, we observed a statistically significant 3-fold increase in CD8 T cells in the LCA under LDF, compared to the contralateral artery control, as we and others have reported previously^41,42^ (Figure 4D&E). In ECKO mice, however, the expected increase of CD8 T cells in LDF vessels compared to contralateral normal laminar flow vessels did not occur (Figure 4D&E, P<0.01). This reduction in CD8 T cell accumulation was similar across male and female ECKO mice (SI Figure 13), and notably did not always coincide with substantial reductions in other immune cell subsets in plaque (CD45 and CD11b, Figure 4D&E). Similarly, this difference in plaque CD8 T cells does not reflect a lack of circulating cells, as there were no differences in CD8 T cells in either blood or lymph node (SI Figure 13). Thus, CD8 T cell accumulation in plaque was strongly suppressed by the deletion of endothelial Elavl1 in the PCAL model.

Next, we tested whether CD8 accumulation was dictated by plaque size, or if it was independent of plaque development. To address this, we examined aortic arches of hypercholesteremic mice at 3 months of age, where we found plaque size was not substantially affected by ECKO (Figure 4G&H). Again, despite similar levels of CD45 and CD11b cells, we found mean levels of CD8 T cells were reduced by ∼70% (Figure 4I&J, P<0.01). Additionally, this ∼70% reduction in CD8 T cells was consistent across sexes (SI Figure 13). To gain a deeper understanding of all of the immune subtypes affected in the plaque, we applied CITE-seq single-cell analysis, with a panel of 103 immune-related surface antigens to examine immune cell composition and expression in the blood and whole aortas of 9-month-old chronic hypercholesterolemic mice. Samples from ECKO and WT mice were pooled in a hash-tagged reaction (N=3 male and N=3 female from each genotype). Cell clustering was performed, using both RNA and protein markers to annotate (Figure 4K&L, SI Figure 14,15). In blood, we again found that overall levels of T cells, monocytes, and granulocytes were very similar between WT and ECKO (SI Figure 14). In contrast, immune cell numbers in aortic tissue were sharply reduced (Figure 4M, SI Figure 14). The major portion of these cells were the CD8 effector memory cells (CD8 TEM), which exhibited a ∼60% reduction that closely approximated the overall reduction we had seen by flow cytometry (Figure 4M, SI Figure 15). Closer examination of RNA and protein markers on CD8 TEM cells within the plaque indicated that the largest differences in RNA and protein expression were associated with TCR signaling, as CD3g and Tcrg-C4 mRNA were reduced (Figure 4N), along with reduced protein levels of TCR and CD38 (Figure 4O). Thus, accumulation of activated CD8 T cells is diminished in plaque tissues of ECKO mice — despite little discernable effect on the circulating CD8 T cell population.

### Endothelial Elavl1 is a stromal checkpoint for CD8 T cell persistence in plaque

Plaque phenotypes suggested that the deletion of Elavl1 from the endothelium inhibits T cell accumulation, and transcriptional data suggests that this may be due to effects on antigen response and activation, rather than recruitment. Activation of CD8 T cells *in vivo* and analysis of their accumulation in atherosclerotic plaque in the PCAL model following our published protocol^42^ confirmed that there was no deficit in activated T cell recruitment to lesions (SI Figure 16). Therefore, to assess effects on T cell persistence after antigen stimulation, we developed an *in vitro* system, co-culturing the cell types necessary to recapitulate the phenotype seen *in vivo*: ECs, antigen-presenting bone marrow-derived dendritic cells (BMDCs), and CD8 T cells. We generated ROSA-CreERT2 ECs with floxed Elavl1 to allow us to excise Elavl1 *in vitro* by treatment with tamoxifen metabolite 4-hydrotamoxifen (4OHT). As atherosclerotic plaque antigens are not yet well defined, we used genetically engineered OT-I CD8 T cells specific to processed ovalbumin (Ova)-derived peptide presented via MHC-I on BMDCs. Binding of the OT-I CD8 T cell receptor to MHC-I-OVA peptide on the BMDC (Signal 1) activates T cell expansion and survival, provided sufficient co-stimulatory (Signal 2) and survival cues (Signal 3) are present. CD8 T cell were marked by CD45.1 and APCs with CD45.2, allowing us to separate these cells from each other and the co-cultured CD45 negative ECs by flow-cytometry markers (Figure 5A&B). In this system, we tested the effect of ECs with and without deletion of Elavl1 on T cell persistence.

T cell response to antigen typically peaks between 7-15 days^43^ so T cell proliferation and survival were examined after 13 days in co-culture. As expected, there was a marked increase in CD8 T cell number (∼2.5 fold, P<0.001), in response to full-length antigen Ovalbumin when co-cultured with same-sex WT ECs and BMDCs (Figure 5B&C). However, deletion of Elavl1 in ECs entirely blocked this proliferative response, despite the same number of WT BMDCs in the system available to present antigen (Figure 5B&C). This was not a result of a generally impaired response to cytokine, since pulsing of BMDCs with already processed SIINFEKL peptide prior to CD8 T cell addition elicited similar responses on both EC types (Figure 5D). To examine the kinetics, we collected co-cultured cells after three days, and found a mild but significant increase in CD8 T cell counts on WT ECs and a lesser and non-significant increase on ECKO ECs (Figure 5E&F). However, the largest observable defect was in the viability of T cells on ECKO cells, which sharply declined (Figure 5E&F). Again, this was not due to a significant difference in the BMDCs (Figure 5E&F, and SI Figure 17). Thus, the data suggest that despite an equivalent ability to respond to an unprocessed antigen (SIINFEKL), CD8 T cells cultured on ECKO ECs are impaired in their survival and persistence in response to processed antigen (ovalbumin).

Our *in vivo* data had suggested that deletion of Elavl1 in ECs affected the translation of several transcripts involved in immune-endothelial interactions. One of the most prominent is C1q. Prior work had shown that C1q suppresses CD8 T cell proliferation in response to low levels of antigen stimulation, but not strong activation by SIINFEKL^44^ – phenotypes which resembled the effects of Elavl1 deletion *in vitro*.

Therefore, we assessed C1q expression in WT and ECKO ECs in culture. We found that, in baseline culture conditions, C1qa–c were not expressed in ECs. However, after co-culture with CD8 T cells, that C1qa could be detected in sorted Icam2+/CD45neg ECs (Figure 5F). Notably, expression could be observed in both WT and ECKO ECs, consistent with prior data showing that the transcript is increased in regions of LDF, even in WT ECs^28^. However, despite the mRNA expression of C1qa, very little C1q protein could be detected in the media when co-culture was performed with WT ECs. In contrast, we observed a substantial increase in C1q protein in the media from ECKO co-culture, consistent with increased levels of translation of C1q in ECKO ECs, and the increase in TRAP associated C1qa–c we had observed *in vivo*. Together, our data shows that Elavl1 deletion from ECs limits the expansion of wild-type CD8 T Cells, even in the presence of their cognate antigen and wild-type antigen presenting cells. These findings suggest Elavl1 acts as a checkpoint in atherosclerosis-derived ECs to suppress CD8 T cell persistence even in the presence of professional antigen presenting cells and cognate antigen.

## Discussion

Here, we demonstrate that Elavl1, which is upregulated in endothelial cells in the atherogenic setting of disturbed flow, suppresses a switch in the ribosomal association of mRNA and T cell persistence in the artery wall. Mechanistically, this includes C1q, a critical regulator of autoimmune responses. These changes can be recapitulated in a novel *in vitro* co-culture system of ECs, bone marrow-derived antigen presenting cells, and CD8 T cells, where reduced Elavl1 in ECs increases C1q protein and reduces T cell persistence following antigen stimulation, even in the presence of wild-type professional antigen presenting cells. *In vivo*, endothelial-specific deletion of Elavl1 reduces CD8 T cell numbers and markers of activation in atherosclerotic lesions without substantial effects on circulating T cell numbers. Together, our data implicate the immune-regulatory transition of the endothelial translatome, marked specifically by C1q, as a crucial determinant of the strength of CD8 T cell responses in atherosclerotic lesions, and the RNA-binding protein Elavl1 as a limiting gatekeeper of this transition. Thus, Elavl1 low endothelial cells are capable of actively repressing immune responses, mimicking properties of regulatory T cells and tolerizing dendritic cells^45^.

### Endothelial Elavl1 and the persistence of CD8 effector memory cells in plaque

Our data show that Elavl1 is required for the accumulation of activated CD8 T cells atherosclerotic plaque. The specific antigens in atherosclerotic lesions are likely diverse; there is evidence that T cells in plaque can be activated by viral infections, oxidized apolipoproteins, and autoimmune disease^21^. Regardless of their initial stimulation, once recruited to plaque, CD8 T cells persist for months at least as effector memory cells (CD8 TEM)^18,42^. Consistent with enrichment of CD8 T cells with antigen-specific response, human coronary arteries have shown high clonality in plaque CD8 T cells, which increased according to the stage of the plaque^46^. Our findings show how the stroma, and the endothelium in particular, contributes to the local maintenance of CD8 TEM. Since Elavl1 expression is potently induced by platelet interactions with the arterial wall, it suggests that Elavl1 induction in the endothelium may be an important checkpoint towards T cell persistence in plaque. As a stromal checkpoint, this would allow for local modulation of immune responses, as we show here – through effects on plaque recruited but not circulating levels of CD8 T cells. Approaches targeting local checkpoints are likely to provide important advantages for local immune-suppression without many of the consequences of systemic immunosuppression.

Mechanistically, our data suggest increased translation of C1q as one possible mediator of the difference in persistence. Ribosomal association of C1q is increased in Elavl1-deleted endothelial cells *in vivo*, and protein production is increased in our *in vitro* co-culture system. Genetic C1q deficiency causes systemic lupus erythematosus (SLE)^47^, which carries a ∼10-20-fold elevated risk for cardiovascular disease^48,49^. Conversely, variants in C1q linked to increased expression are protective against autoimmune disease^50^. In an animal model, loss of C1q increased plaque development^51^. While C1q is important in the opsonization of dead cells, treatment of CD8 T cells in culture with C1q directly limits antigen-triggered expansion and activation^52^. Thus, our data provide a cellular and post-transcriptional mechanism regulating a recently described transition in the arterial endothelium in atherogenic conditions, termed an immune-cell like phenotype (EndICLT), which is linked to the expression of C1q^28^. Furthermore, we show that this transition plays an important role in the persistence of activated T cells, suggesting that the upregulation of Elavl1 in the endothelium is an important contributor to the breakdown of local barriers to autoimmunity observed in atherosclerotic lesions^53^.

### Translational regulation in atherosclerosis

Here, we show that Elavl1 limits immunosuppressive changes in mRNA translation in the atherogenic endothelium. Translation of mRNA is a tightly regulated process, and variations between mRNA isoforms can lead to >100-fold differences in protein translation^54^. However, very little is known about the regulation of translation during chronic endothelial activation in atherosclerotic lesions. In myeloid cells and the endothelium, Elavl1 has principally been considered in the context of 3’UTR binding and transcript stability^55,56^. Given this emphasis in the literature, it was somewhat surprising to us that the major effect we observed was not on canonical Elavl1 stabilized transcripts. Such analysis may require additional *in vivo* experiments to examine transcript kinetics. Nevertheless, whatever these effects may be, they do not appear to strongly affect canonically stabilized transcripts (e.g., Ifit5, Nfkb1, Nfkb2, Ifitm3, Irf9, Mapk1, Ifnb1, Ptgs2/Cox-2, Tgfb, Ctss, Sele)^55,57^ in either the total mRNA or TRAP mRNA pool. The finding that Elavl1 can instead regulate ribosomal binding and protein translation is new in this context but has been reported in epithelial cells and B cells. In conditions of hypoxia in epithelial cells, an unbiased proteomic analysis revealed a 3-4 fold enrichment of Elavl1 (and also Ptbp1) at the polysome^58^. Previous analysis of the effect of Elavl1 loss in the endothelium had revealed alterations in the splicing of the transporter of a key translational protein, eIF4e transporter (Eif4enif1), which we confirmed^59^. In future work, it will be interesting to examine whether Elavl1 effects on T cell responses are coupled with cell metabolic alterations.

Supporting this idea, Hif2a enhanced polysome association of Elavl1 in epithelial cells^58^, and Hif2a levels are increased in ECs exposed to disturbed flow in the absence of hypoxia^60^. Thus, changes in ribosomal association of specific mRNA and the RBPs which regulate this process may be essential in the coupling of metabolic dysfunction and the immune response.

Together, our work shows that changes in the translatome in atheroprone endothelium is concentrated in genes involved in the regulation of chronic immune-endothelial interactions, defining new targets to modulate chronic immune-endothelial interactions. The RNA-binding protein Elavl1 limits an immunosuppressive component of this response, functioning as a novel stromal checkpoint to T cell persistence in atherosclerotic lesions. We predict that this may be important not only in the accumulation of CD8 effector cells in plaque, as we show here, but also likely in other settings of chronic immune responses, including autoimmune disease and graft-vs-host responses, and in settings where effector cell accumulation is impaired, as in cancer immunotherapy and anti-viral responses in old age. The endothelium, found in all vertebrates, evolutionarily predates the appearance of T cells in jawed vertebrates. Given this, it is not surprising that as the adaptive immune system developed, it was entwined with endothelium, which transports immune cells into tissues. Therefore, we expect that this is just the beginning of work to understand how these interactions locally modify immune cell function.

## Materials and methods Mice

For Elavl1 EC knockout (KO) experiments, *Cdh5(PAC)-CreERT2* and *Elavl1f/f* mice were used and have been previously described^61,62^. They were inter-crossed to generate Elavl1 ECKO mice (*Cdh5(PAC)- CreERT2; Elavl1f/f*) and littermate controls (*Elavl1f/f*) used here. For PCAL and 3-month chronic hyperlipidemia models, Elavl1 excision was accomplished by intraperitoneal injections of either 1 mg (100µL of 10mg/mL) Tamoxifen (Sigma T5648) in sunflower oil three times, every other day, at least one week prior to experimental perturbations. Mice were used for experiments in sex-matched and litter-matched pairs.

For TRAP RNAseq experiments, the *Rosa26-fsTRAP* have been previously described^29^. *Rosa26-fsTRAP* mice were inter-crossed with *Cdh5(PAC)-CreERT2; Elavl1f/f* mice to generate Elavl1 ECKO RiboTRAP mice *(Cdh5(PAC)-CreERT2; Elavl1f/f; Rosa26-fsTRAP)*. To generate Elavl1 WT RiboTRAP control mice (*Cdh5(PAC)-CreERT2; Rosa26-fsTRAP)*, *Rosa26-fsTRAP* mice were inter-crossed with *Cdh5(PAC)- CreERT2.* Tamoxifen was delivered as above.

For generation of immortalized Elavl1 ECKO mAECs, previously described *R26-CreERT2* (G*t(ROSA)26Sor^tm^*^1^*^(cre/ERT2)Tyj^*/J) mice^63^ were inter-crossed with *Elavl1f/f* mice to generate *RosaCreER; Elavl1f/f* experimental mice.

For OT.1 T cell isolation, Ovalbumin (SIINFEKL257-264)-specific OT.1 TCR transgenic recombination activation gene-1 deficient (*Rag1*^-/-^) on C57BL/6J mice background were used. Mice were homozygous for CD45.1 or as previously described^64^. Wild-type (CD45.2) bone marrow cells were isolated from C57BL/6J mice.

All mice were housed and handled in accordance with protocols approved by the University of Connecticut Health Center for Comparative Medicine.

### Models of Atherosclerosis

For both chronic and PCAL models, hyperlipidemia was induced via 100µl intraperitoneal injection containing 1E11 viral particles of AAV8-mPCSK9 (pAAV/D377Y-mPCSK9) (Roche-Molina, ATVB, 2015) produced at the Gene Transfer Vector Core (Schepens Eye Research Institute and Massachusetts Eye and Ear Infirmary, Harvard Medical School). Mice were also placed on Clinton/Cybulsky high-fat rodent diet (HFD) with regular casein and 1.25% added cholesterol (Research Diets, D12108C). Hyperlipidemia was confirmed after blood collection via cardiac puncture through the right ventricle. Serum was collected after centrifugation and stored at -80°C until analysis by Total Cholesterol Assay Kit (Cell Bio Labs INC, STA- 384).

Partial carotid ligation surgery (PCAL) was performed as previously described^22^. Briefly, while under isoflurane anesthesia, the left external carotid (LCA), internal carotid, and occipital arteries were ligated. Mice were treated post-surgery with Buprenex or Ethiqua as analgesic. High resolution Doppler ultrasound was performed to confirm flow reduction in the LCA.

### Total Carotid Intimal EC RNA Collection

Elavl1 ECKO mice (*Cdh5(PAC)-CreERT2; Elavl1f/f*) and littermate controls (*Elavl1f/f*) were treated with tamoxifen (3x 1mg, intraperitoneal) and PCAL and hyperlipidemia induced as described. Total carotid intimal RNA was isolated 3 weeks after PCAL induction. Carotid arteries were dissected and rapidly flushed with 150uL of Trizol (Thermo, 15596026). Trizol eluant was stored at -80°C until the time of RNA isolation. For RNA isolation, chloroform was added at 1:5 ratio to Trizol (30uL:150uL Trizol) and the clear supernatant containing mRNA was combined 1:1 with 70% Ethanol (EtOH) and then ran on Qiagen RNeasy Micro Kit (Qiagen, 74004).

### TRAP RNA Collection

Elavl1 ECKO RiboTRAP mice (*Cdh5(PAC)-CreERT2; Elavl1f/f; Rosa26-fsTRAP*) and controls (*Cdh5(PAC)- CreERT2; Rosa26-fsTRAP*) were treated with tamoxifen for recombination and then subjected to PCAL and hyperlipidemia, as described above. Carotid TRAP RNA was isolated 3 weeks after PCAL induction. Mice were perfused through the left ventricle with cold PBS. Carotid arteries were dissected and flushed with cold Lysis Buffer (20mM RNAse free HEPES Potassium Hydroxide pH7.4, 5mM RNAse free Magnesium Chloride, 150mM RNAse free Potassium Chloride, 0.5mM Dithiothreitol, Complete protease inhibitor EDTA free, 100µg/mL Cycloheximide in Methanol, 40U/mL RNAse inhibitor, 1% NP-40, 30 mM 1,2-diheptanoyl-sn-glycero-3-phosphocholine (DHPC) in RNAse free PBS) to extract intimal protein and RNA.

GFP- (HtzGFP_04, clones 19C8 and clone19F7, Memorial Sloan Kettering Antibody and Bioresource Core) or IgG- (Sigma, 15006) conjugated M270 Dynabeads (Fisher, 14301) were prepared per manufacturer’s instructions. Lysate was then incubated with 1 mg of conjugated beads per vessel at 4°C for 30 min. After immunoprecipitation for 1 min at 4°C, supernatant was collected and combined with RNA lysis buffer RLT (Qiagen) plus Beta-mercaptoethanol. The bead-bound fraction of RNA and protein was washed three times with cold Potassium Chloride wash buffer (20mM RNAse free HEPES Potassium Hydroxide pH7.4, 5mM RNAse free Magnesium Chloride, 350mM RNAse free Potassium Chloride, 1% NP-40, 0.5mM Dithiothreitol, 100µg/mL Cycloheximide in Methanol). Beads were then resuspended in RNA lysis buffer RLT (Qiagen) plus Betamercaptoethanol. All samples were then stored at -80°C.

### RNA isolation and preparation for qPCR and RNA Sequencing

For qPCR analysis, cDNA was generated using the High-Capacity cDNA Reverse Transcription Kit (Fisher, 4368814) per the manufacturer’s instructions. qPCR reactions were generated using iQ SYBRGreen Supermix (Bio-Rad, 1708880) and run on Bio-Rad CFX96. For total intimal-RNAseq, samples were prepared with Illumina TruSeq stranded RNA library and sequenced at 25 million reads per sample, 75bp single end reads. For TRAP-RNA sequencing, samples were prepared using Takara Bio v4 SMARTSeq (Takara, 635026) and sequenced at 100 million reads per sample, 150 bp paired-end sequencing.

### Gene Expression and Alternative Splicing Analysis

FASTQ files were aligned via STAR against reference genome Gencode M25 (GRCm38.p6) and gene expression was quantified by RSEM. Count tables were then analyzed by DESeq2^65^ in R. Expression levels were examined by TpM. Intron clustering, requiring 50 split reads supporting each cluster, and differential splice analysis were performed via Leafcutter^66^. Subsequent data analysis to examine pathway enrichment was performed in Jupyter Notebook, with code outlined here (https://github.com/pamurphyUCONN/2024_Nicholas).

### *En Face* Immunofluorescence

Mice were perfused through the left ventricle first with cold phosphate buffered saline (PBS) and then cold 0.2% paraformaldehyde (PFA) in PBS. Aortas were dissected, visceral fat was removed, and the vessel was flushed again with PBS. Vessels were then pinned to wax and dissected to expose the arterial intimal lining. Vessels were fixed in 0.2% PFA for 1 hour at 4°C, followed by blocking and permeabilization in 1% bovine serum albumin (BSA) in PBS containing 0.2% Triton X-100 for 1 hour at room temperature (RT). Vessels were incubated in primary antibody overnight at 4°C in 0.1% BSA with 0.02% Triton X-100. Primary antibodies used were rat anti-mouse VE-Cadherin at 1 µg/mL (BD, 550548) and rabbit anti-mouse Elavl1 at 7 µg/mL (Abcam, 200342). After three washes with cold PBS, vessels were incubated with secondary antibodies for 1 hour at RT in 0.1% BSA with 0.02% Triton X-100. Secondary antibodies used were goat anti-rat AF488 at 2 µg/mL (Thermo, A11006) and goat anti-rabbit AF594 at 2 µg/mL (Thermo, A11037). Vessels were then washed for 5 minutes at RT with PBS and 1 µg/mL DAPI (Sigma, D9542). Two washes with PBS followed before mounting on a slide with Fluoromount (Sigma, F4680) for imaging on LSM800 confocal microscope with ZenPro software. After image collection with the same instrument settings and during the same session, Mean Fluorescence Intensity (MFI) was calculated using ImageJ.

### Atherosclerotic Plaque Severity Analysis

For both chronic and PCAL models, blood was collected via cardiac puncture through the right ventricle. Mice were then perfused through the left ventricle with cold PBS. Arteries were dissected, adventitia and visceral fat was removed, and the vessel was flushed again with PBS. For chronic hyperlipidemia, plaque in some aortas was aortas was directly imaged in most cases, by opening the artery lengthwise, pinning it down on black wax and taking images on a dissection scope with standardized magnification. Plaque area was quantified using ImageJ by combining all images and thresholding together to select plaque ROIs.

For partial carotid ligation analysis, the midpoints (containing about ∼1/3 total tissue) from carotid arteries were fixed in 4% PFA overnight at 4°C, then incubated in 30% sucrose in PBS for several hours before embedding in OCT (Fisher, 23730571). Carotid arteries were then snap-frozen and stored at -80°C until cryo-sectioning at 10μm. Oil Red O staining was performed by incubating tissues with a 0.22 filtered 3:2 dilution of 6.25g/L Oil Red O in isopropanol to distilled water for 5 min before washing with tap water. Slides were mounted and then imaged. Percent occlusion was quantified using ImageJ.

### Plaque immunophenotyping

#### Vessel and Lymph Node Digestion

Aortic vessels were obtained after PBS perfusion and visceral fat and adventitia were removed. Vessels were mechanically digested via mincing for 4 min each and then enzymatically digested for 1hr at 37°C with gentle rotation. Mesenteric and cervical lymph nodes were isolated and minced similarly, before enzymatic digestion. Enzymatic digestion cocktail was prepared in balanced salt solution (BSS) with 150 U/ml collagenase type IV (Sigma-Aldrich C5138), 60 U/ml DNase I (Sigma-Aldrich), 1 µM MgCl_2_ (Sigma Aldrich), 1 µM CaCl_2_ (Sigma-Aldrich), and 5% fetal bovine serum (FBS). Digested tissue was gently crushed over 35µm nylon mesh cell filter caps and digestion was quenched with 10% FBS in BSS. Samples were centrifuged for 5 min at 300 x g at 4°C, supernatant was aspirated, and the cell pellet was resuspended in flow cytometry buffer (FACS buffer), 2% FBS and 1 mM Ethylenediaminetetraacetic acid (EDTA) in PBS.

#### Blood collection

Blood was collected via cardiac puncture at the time of collection. An aliquot was reserved for serum cholesterol analysis. The remainder was immediately added to 0.55g/L heparin (Sigma, H3393-50KU) in HBSS and temporarily stored on ice. Samples were were centrifuged for 5 min at 300 xg. Supernatant was aspirated and pellets were resuspended in Cold Spring Harbor Red Blood Cell Lysis (155 mM NH4Cl, 12 mM NaHCO3, 0.1 mM EDTA) and incubated at RT for 10 minutes then quenched with PBS. Lysis was repeated if needed. Samples were centrifuged for 5 min at 300 xg at 4C, supernatant was aspirated, and the cell pellet was resuspended in FACS buffer.

#### Immunostaining

Cell pellets were resuspended in the following antibody cocktail diluted in FACS buffer: 1 µL per test Fc Block (InVivoMAb anti-mouse CD16/32, BioXcell BE0307), LIVE/DEAD UV Blue (1:100, ThermoFisher, L34962), CD8 (1:200, Biolegend, 100708, clone 53-6.7), CD4 (1:200, Biolegend 100536, clone RM4–5), CD45.2 (1:200, Biolegend, 103116, clone 30-F1111), Cd3e (1:200, Biolegend, 100348, clone 145-2C11), Gr1 (1:200, B.D. Pharmingen, 552093, clone RB6-8C5), and Cd11b (1:200, Biolegend, 10112, clone M1/70). Full-minus one (FMO) controls were used to set gates. LSRIIA (BD Biosciences) was used for acquisition. All flow cytometry data were analyzed with FlowJo (Tree Star, Ashland, OR).

#### 10X Single Cell CITE-Seq for chronic hyperlipidemic samples

Whole and PBS perfused aortic tissues were similarly dissected of adventitia followed by a slightly modified digestion protocol. Vessels were minced for 4 minutes per sample and then enzymatically digested for 37C at 265 RPM for 50 minutes. Digestion cocktail was prepared in HBSS (Fisher, 24020117) with Collagenase Type I (Worthington, CLS-1) 1000 U/mL, Collagenase Type XI (Sigma, C7657) 180 U/mL, Hyaluronidase Type I-s (Sigma, H3506) 80 U/mL, Dnase I (Sigma, DN25) 80 U/mL, and FBS (1:200). Digestion was quenched with 100% FBS before tissue was gently crushed over 100 um nylon mesh cell filters. Samples were centrifuged for 5 min at 300 xg at 4C, supernatant was aspirated, and the cell pellet was resuspended in FACS buffer.

Aortic samples were stained for flow cytometry in FACS Buffer with 1 uL per test Fc Block (InVivoMAb anti-mouse CD16/32, BioXcell BE0307), anti-mouse CD45 (1:200, Biolegend, 146603, clone I3/2.3), CD31 (1:200, Biolegend, 102509, clone MEC13.3) and ICAM2 (1:200, Invitrogen, A15451, clone 3C4). To hashtag samples for 10x reaction, cells from each tissue were stained with a unique TotalSeqB anti-mouse hashtag (1:200, Biolegend) and incubated for 30 minutes at 4C. DAPI was added to aortic samples immediately before FACS to distinguish live cells. CD45+ live immune cells from all aortic samples were collected into one tube and CD45-CD31+ICAM2+ live endothelial cells from all aortic samples were collected into another protein lo-bind 1.5 mL Eppendorf tube pre-coated with 1% BSA in PBS.

All FACS sorted cells and were then stained with CITE-Seq TotalSeqB mouse Universal Cocktail (Biolegend, 199902) and TotalSeq-B0924 CD257 OX-40L (Biolegend 108825) for 30 minutes at 4C for CITE-Seq analysis. An equal number of white blood cells from each sample were combined and then also stained with this antibody cocktail under the same conditions. After washing, approximately 40,000 cells from blood and 40,000 cells from aorta were then combined into a single 10X reaction (Chromium Next GEM Single Cell 3’ Kit v3.1) and run on a Chromium system to generate single cell cDNA library. Steps followed the 10x protocol, with the addition of a 3min extension during library amplification to reduce amplification bias.

#### Analysis of 10X Single Cell CITE-Seq

Cellranger (7.0.1) was used to map reads against Mouse (mm10) 2020-A and to generate gene level and antibody level read counts. Paths to feature library (antibody capture and gene expression) were specified during the processing, so we ended with a h5 object that included both gene expression and antibody capture information. Raw h5 feature-barcode matrix files were analyzed using the Seurat package (5.0.3). Hashed samples were demultiplexed using the MULTIseqDemux function (autoThresh = TRUE) which is part of the deMULTIplex R package and cells labeled with multiple, or no hashtags were removed. Cells with <300 or >5000 genes, > 20000 UMI, and/or > 10% mitochondrial reads were filtered out. Remaining doublets were filtered out using scDblFinder Bioconductor package (1.18.0). Data from all objects was integrated and normalized using the sctransform R package (0.4.1) and Principal Component Analysis (PCA) was performed using the 3000 most variable genes and “RunPCA” function in the Seuart Package. Clustering was carried out using the first 20 PCs and the “RunUMAP”, “FindNeighbors”, and FindClusters functions in the Seurat package. UMAPs were visualized using the “DimPlot” in the Seurat package.

Clusters were annotated manually by analyzing enriched transcripts. Subclustering was performed by subsetting clusters associated with cell types of interest and performing data integration (sctransform) and UMAP clustering as described above. Differential expression analysis was performed by using the “FindMarkers” function in the Seurat package with the Wilcoxon rank sum test algorithm and differences with and adjusted p-value < 0.05 were considered significant. Antibody barcode counts were normalized using centered log ratio (CLR) normalization and differential expression of normalized antibody counts was performed using the “FindMarkers” function in Seurat as previously mentioned.

### Generation of *in vitro* Endothelial Cell Line

Mouse aortic endothelial cells were isolated from *RosaCreER; Elavl1f/f* adult male mice as previously described^22^. Briefly, the vasculature was perfused with cold PBS through the left ventricle. The descending aorta was dissected and filled with 2% collagenase type II (Worthington, LS004176) in serum-free DMEM then closed at the ends with 7-0 suture. The digesting vessel was incubated for 45 minutes at 37°C in DMEM + 20% FBS. Subsequently, digested intimal cells were flushed out with vascular basal medium + endothelial cell growth kit VEGF (ATCC, PCS-100-030 and PCS-100-041). Flushed cells were centrifuged at 300 x g for 3 minutes, supernatant aspirated, and cells were plated on rat tail collagen type I (Fisher, A1048301) in EC growth medium (as above) supplemented with primocin (1:500, Invivogen, ant-pm-1).

Intimal cells were expanded for 48 hrs and then treated with lentiviral TetOn-SV40 and polybrene (1:1000, Sigma, TR-1003-G). Viral vector was constructed as described^22^. Cells were expanded in EC growth medium and 2 µg/mL doxycycline (dox, Sigma, D9891-1G). For FACS purification, cells were stained from 30 min at 4°C in FACS buffer with CD45-APC-Cy7 (Clone 30-F11, Biolegend 103116) and ICAM2-PE (Clone 3C4, Invitrogen, 63-0441-82) and FACS sorted (BD AriaII) for single-cell CD45-ICAM2+. Sorted aortic ECs were expanded in the presence of dox, immortalized with retroviral pBABE-neo-hTERT (Addgene, 1774), and then allowed to expand. To induce excision of Elavl1 in *RosaCreER; Elavl1f/f* immortalized mAECs prior to coculture experiments, cells were treated with a final concentration of 1µM 4OHT (Z)-4-HydroxyTamoxifen (4OHT, Sigma H7904) for three days or with 100% EtOH daily as vehicle control.

## Western Blot

Media was collected and cells were washed 3x with PBS. Cell lysates were collected in RIPA buffer using cell scrapers and then stored at -20°C. Lysate was thawed, treated with 1M Dithiothreitol (DTT, Thermo, A39255), incubated at 95°C for at least 5 minutes, and then diluted 1:1 with 2X Laemelli buffer. Samples were run on Novex WedgeWell 4-20% Tris-Glycine Mini Gel (Thermo, XP00100BOX) at 125V for 1.5 hrs. Transfer to membrane was done using Trans-Blot Turbo Transfer System (BioRad, 1704150) and transfer kit (BioRad, 1704273).

Membranes were first blocked in 5% milk in tris buffered saline + 0.1% Tween-20 (TBST) for 30 min at RT. Membranes were then incubated in primary antibody for Elavl1 (1:1000, Abcam, 200342) ON at 4°C in 0.5% milk in TBST. After washing in TBST, membranes were incubated in secondary antibody for anti-rabbit-HRP (1:5000, Jackson Immuno, 111-035-144) for 1 hr at RT. After developing and imaging with Clarity ECL Western Blot substrate (Biorad, 1705060), the membrane was washed again. The membrane was then blocked in 5% milk TBST for 30 min at RT, incubated with BetaActin-HRP (1:1000, Invitrogen, MA5-15739-HRP) for 3 hrs at RT. Washing, developing and imaging followed staining as described above. Quantification was performed using ImageJ.

### *In Vitro* Co-Culture System

#### Antigen Presenting Cell Generation

Bone marrow was collected from the femurs and tibia of adult male C57BL/6J mice. Briefly, bones were dissected, ends removed, and placed open end down into 0.5 mL Eppendorf tube with 18G needle puncture. The bones and punctured 0.5ml tube were inserted into a 1.5 mL Eppendorf and then centrifuged at 21.2K x *g* for 15-20 seconds. Pelleted bone marrow was resuspended in PBS, passed over a 100 µm filter and centrifuged again at 300 x *g* for 5 min. Supernatant was decanted and cells were treated with red blood cell (RBC) lysis buffer (155mM ammonium chloride, 12 mM sodium bicarbonate, 0.1 mM EDTA) for 1 min. RBC Lysis was quenched with FACS buffer. Cells were pelleted by centrifugation at 300 x *g* for 5 min, then counted on Luna cell counter (New England BioGroup, L20001). Monocytes were then isolated using EasySep Monocyte Isolation Kit (StemCell Tech, 19861). Monocytes were seeded in immune cell medium (MEM with L-glutamine, 10% FBS + 50 mL Tumor Cocktail, 2 µg/mL primocin) supplemented with 20 ng/mL granulocyte-macrophage colony-stimulating factor (GM-CSF, Biolegend, 576304) on non-tissue culture treated plates. After 3 days of culture, cells were fed with fresh media and 40 ng/mL GMCSF. On day 5, non-adherent and loosely adherent cells were collected, resuspended in APC cryo preservation media (90% immune cell media, 10% dimethylsulfoxide (DMSO), 20 ng/mL GMCSF) and stored at -80°C.

#### CD8^+^ T Cell Isolation

Spleens from OT.1 *Rag^-/-^* transgenic mice were dissected and then crushed over 100 µm cell strainer. Cells were pelleted by centrifugation at 300 x *g* for 5 minutes. Supernatant was decanted and pellets were resuspended in RBC lysis buffer in 1–2mL for 2–3 minutes. Lysis was quenched with PBS or BSS. Lysed cells were then passed over 70 µm cell strainer, pelleted by centrifugation at 300 x *g* for 5 min, then counted on Luna cell counter. CD8 T cells were isolated using EasySep CD8 T cell isolation kit (StemCell, 19853).

Isolated CD8 T cells were pelleted and then resuspended in 90% FBS + 10% DMSO for cryopreservation and stored at -80°C.

#### Co-culture Experimental Design

The *RosaCreER; Elavl1f/f* endothelial cell line (+TetOn-Sv40 and +hTert) were plated in 96-well plates at 1x10^4^ cells per well (Day -4), then treated with 1 uM 4OHT or EtOH daily for three days, starting on the day of plating (D-4 to D-2). The next day (D-1), 4.4 x10^3^ thawed APCs were added per well to EC culture, along with or without antigen, either 0.5 ug/mL EndoGrade Ovalbumin (BioVender, 321000) or 1 ug/mL SIINFEKL (Ovalbumin 257-264; Sigma-Aldrich, S7951). The following day fresh media without antigen was changed for wells treated with SIINFEKL. Subsequently, 2.1 x10^4^ thawed naïve CD8 OT.1 T cells were added to all wells regardless of antigen treatment. For EdU incorporation assay, 1 x10^5^ thawed naïve CD8 T cells were added per well along with 1 uM EdU (Thermo, A10044). For co-cultures collected at D13, 100 uL supernatant was removed and 100 uL of fresh media with or without antigen was replaced.

Co-cultured supernatant and cells were then collected 3 days or 13 days later for assay by western or flow cytometry. At the final collection point, supernatant was collected and stored at -20C or -80C until use. For flow cytometry, non-adherent cells (T cells and the majority of APCs) were first removed from all wells. Then adherent cells (ECs and few APCs) were collected via 0.25% Trypsin in Versene for 5-10 minutes at 37C. Trypsin was quenched with FBS and cells were then processed for flow cytometry.

#### Flow Cytometry Staining

CD45.1+ OT.1 co-cultured cells were stained with an antibody cocktail diluted in FACS buffer: 1 µL per test Fc Block (InVivoMAb anti-mouse CD16/32, BioXcell BE0307), LIVE/DEAD UV Blue (1:100, ThermoFisher, L34962), CD45.1 (1:200, Thermo, 25-0453-82, clone A20), CD45.2 (1:200, Biolegend, 109806, clone 104), CD8 (1:200, Biolegend, 100712, clone 53-6.7), CD44 (1:200, Invitrogen, 63-0441-82, clone IM7), and ICAM2 (1:200, Invitrogen, A15451, clone 3C4). BioRad ZE5 was used for acquisition. All flow cytometry data were analyzed with FlowJo (Tree Star, Ashland, OR). Flow cytometry results were analyzed for CD8 T cell count, viability, and proliferation, along with APC count and viability, and endothelial cell count and viability. One biological experimental replicate of D3 experiments in the was excluded based on the lack of APCs recovered at the time of collection in the full-length ovalbumin treated wells. Biological experimental replicates of D13 collection were excluded based on the following criteria: reduced APC viability across all conditions compared to the majority of experimental replicates; statistical outliers, and use of female CD8 T cells as compared to male T cells. The replicate using female T cells had a trend towards fewer T cells detected in the no antigen condition as compared to the other replicates. Endothelial cells and APCs were both derived from male mice.

### Adoptive Transfer for Analysis of Recruitment to Low Flow Carotid Artery

CD8 OT.1 T cells were isolated as above. Elavl1 ECKO and WT mice underwent PCAL and induction of hyperlipidemia. Two weeks later, 2x10^5^ naïve CD8 OT.1 cells were intravenously transferred into PCAL recipient mice. Within 24 hours of transfer, recipient mice received intraperitoneal injection of 50 µg SIINFEKL^257–264^ peptide (Sigma, S7951-1MG), 20 µg of agonist anti-OX40 (clone OX-86 mAB, BioXcell, BE0031) and 10 µg of agonist anti-41BB (clone 3H3 mAb, BioXcell, BE0239-1MG). One week post transfer, carotid arteries were collected, digested as described above, and assayed via flow cytometry. Digested cells were stained with the following antibody cocktail diluted in FACS buffer: 1 µL per test Fc Block (InVivoMAb anti-mouse CD16/32, BioXcell BE0307), Live/Dead Zombie Aqua (1:100, Biolegend, 423101), CD45.1 (1:200, Thermo, 25-0453-82, clone A20), CD45.2 (1:200, Biolegend, 109823, clone 104), CD3e (1:200, Biolegend, 100348, clone 145-2C11), CD4 (1:200, Biolegend, 100536, clone RM4-5), CD8 (1:200,

Biolegend 100708, clone 53-6.7), and Va2 (1:200, Invitrogen, **46-5812-80**, clone B20.1) and Vb5 (1:200, Biolegend, 139503, clone MR9-4). LSRIIA (BD Biosciences) was used for acquisition. All flow cytometry data were analyzed with FlowJo (Tree Star, Ashland, OR).

## Analysis of Public Single Cell Data

Data was accessed via NCBI GEO (https://www.ncbi.nlm.nih.gov/geo/) from SRR13932927-30. Cellranger/6.0.2. in 10x Cloud was used to map reads against Mouse (mm10) 2020-A. Hd5 feature-barcode matrix files were analyzed in Scanpy^67^. First, cells were filtered to meet minimal gene expression (200 genes), then cells with >10% mitochondrial reads were filtered out. Cells with <2000 or >60000 UMI were filtered out. The 2000 most variable genes were used for Leiden clustering. Clusters were annotated manually by enriched transcripts. Differential expression analysis was performed by tl.rank_genes_groups in Scanpy, with Wilcoxon analysis and Benjamini-Hochberg correction.

### Statistical Analysis

With the exception of single-cell analysis (statistical comparisons performed in Scanpy), statistical analysis was performed in GraphPad Prism. For paired samples with equal variance, a Student’s t test was performed, otherwise a Mann-Whitney U test was performed. For comparisons among multiple samples, we used ANOVA with post-hoc Sidak’s multiple comparisons test when variances were equal and Kruskal-Wallis with post-hoc Dunn’s when they were not.

## Data availability

Bulk RNA sequencing data and CITE-seq data is deposited in SRA (PRJNA1142064)

## Supporting information

SI_Table1_MergedDESeq2

SI_Table2_MergedLeafcutter

SI_Table3_RSEM_TpM

## Acknowledgements

We thank Bernard Cook, Science Writer, Editor, and Illustrator for the School of Medicine at UConn Health, for editing and help with our graphical abstract. In the course of this work, we received valuable input from the ImmunoCardiovascular Group at UConn Health. This work was supported by UConn Health startup funds from the School of Medicine and Department of Cell Biology, Center for Vascular Biology and Calhoun Cardiology Center, American Heart Association Innovative Project Award 19IPLOI34770151 (to P.A.M.); NIH National Heart, Lung, and Blood Institute Grants K99/R00-HL125727 and R01-HL150362 (to P.A.M); NIH National Heart, Lung, and Blood Institute Grant R01HL172239 and National Institute of Diabetes and Digestive and Kidney Disease Grant R01DK121805 (to B.Z.); Linda and David Roth Cardiovascular Research endowment (to A.R.); supported in part by the Boehringer Ingelheim endowed chair (to A.T.V.); and American Heart Association Predoctoral award (to S.-A.E.N.).

---

**SI Figure 1.**
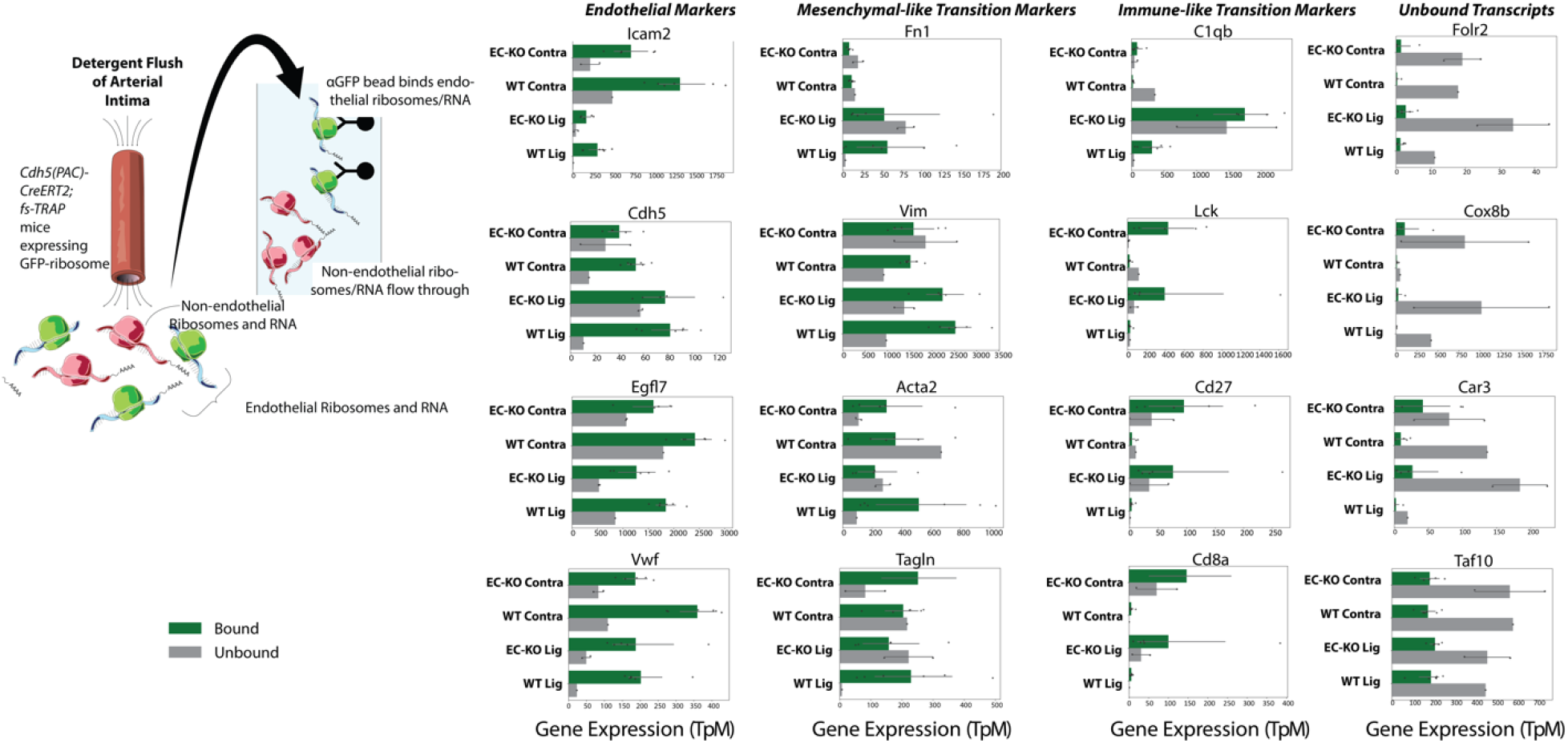
Transcript enrichment by ribosomal pulldown (TRAP) from intimal flush. Figure shows the enrichment of individual transcripts over unbound mRNA in the intimal flush. Each point indicates a biological replicate, from the indicated genotype and in vivo flow profile, either from the low flow (LDF) or contralateral normal flow (Contra) artery.

**SI Figure 2.**
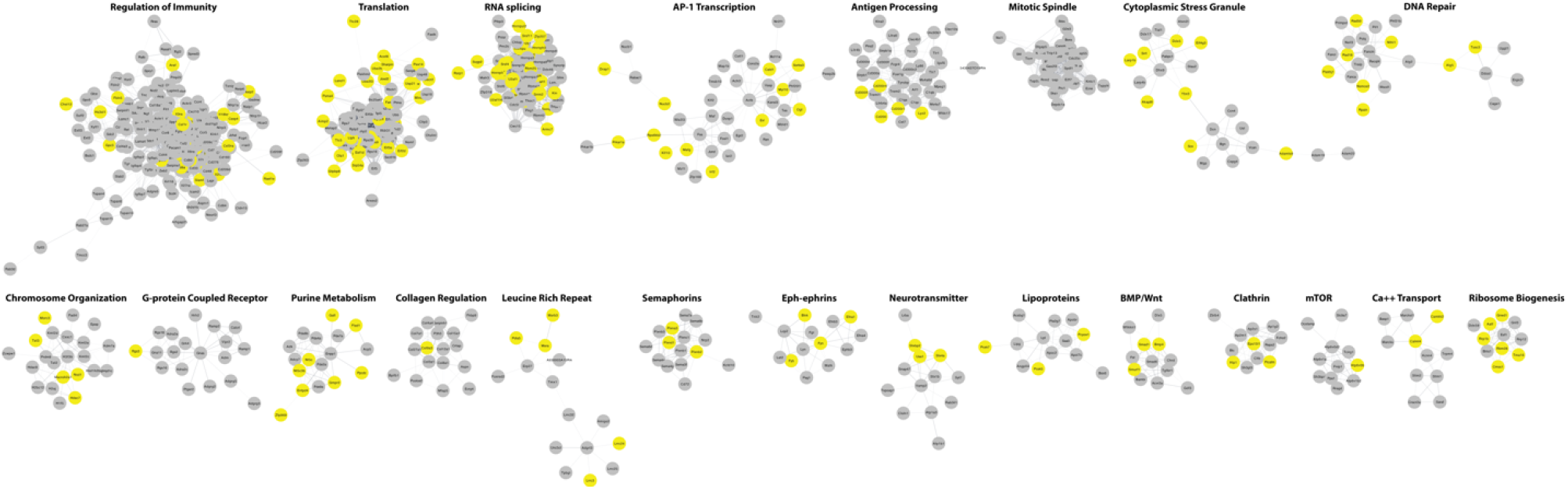
Atherosclerosis-associated transcriptional changes in TRAP-mRNA Cytoscape visualization of all terms associated with low-flow-regulated transcripts in ribosomal mRNA, either by mRNA levels (DESeq2) or splicing (Leafcutter) in TRAP mRNA from endothelial cells in atherosclerotic artery. Expression (gray; padj <0.05) or splicing (yellow; padj <0.05).

**SI Figure 3.**
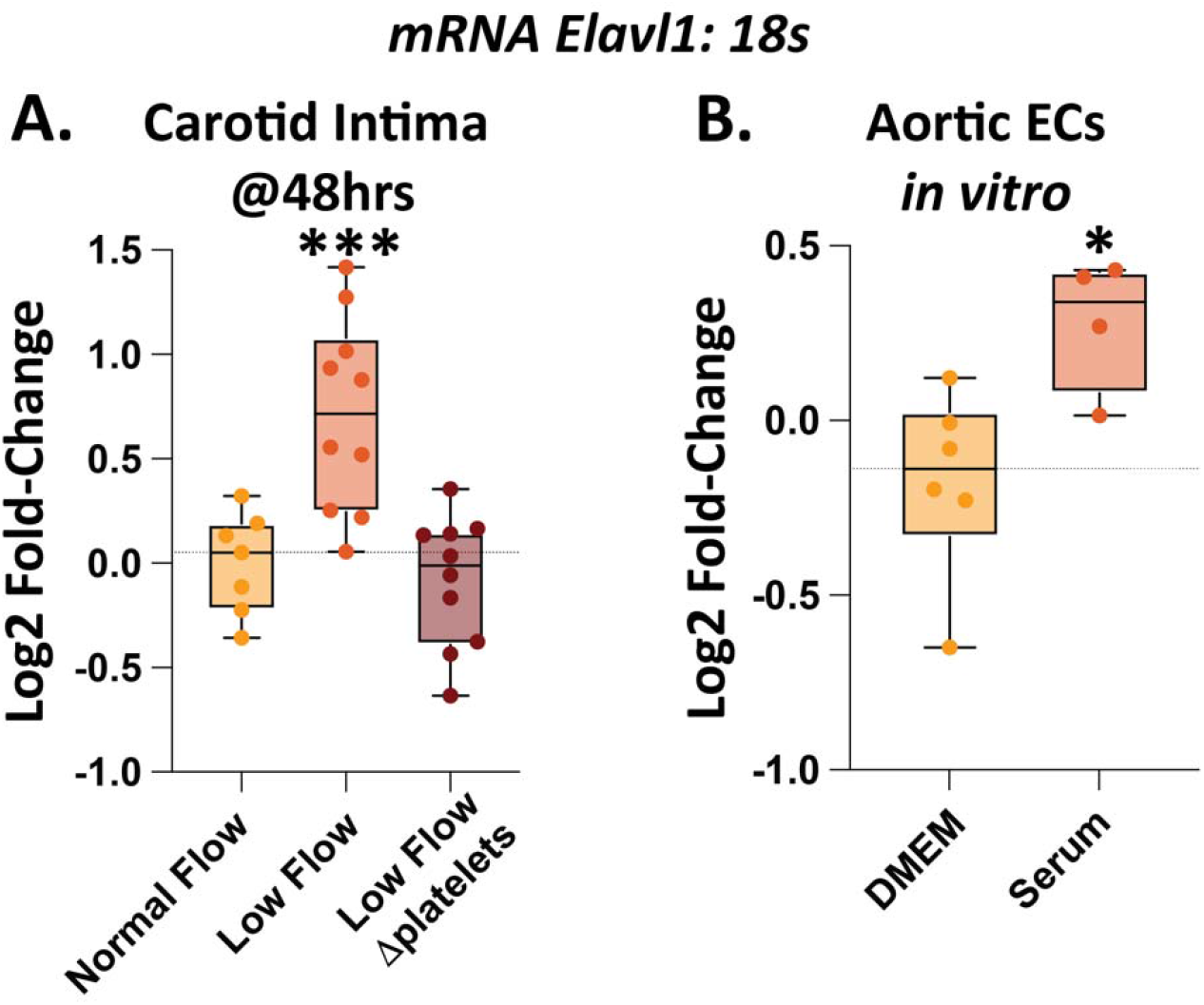
Acute changes in Elavl1 expression under atherogenic conditions are platelet dependent. (A) Log2 Fold Change of Elavl1 mRNA expression as detected by qPCR, normalized to housekeeping gene 18s. (B) Isolation of RNA from mouse aortic ECs in culture in the presence or absence of mouse serum, containing platelet releasate. Each point represents one biological replicate well. ***P<0.001, and *P<0.05. ANOVA with post-hoc Dunnett’s test, or T-test.

**SI Figure 4.**
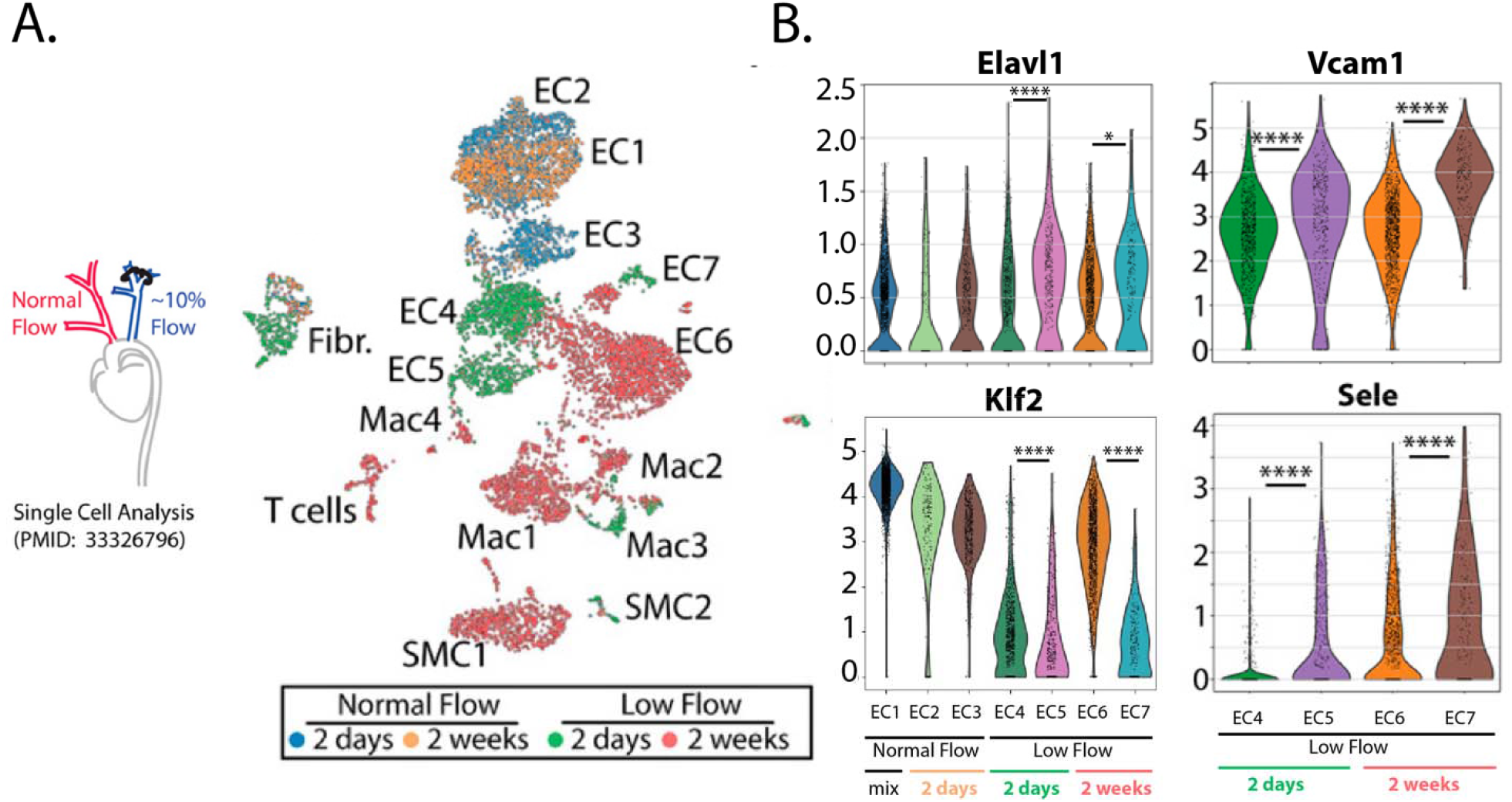
Low and disturbed flow-induced Elavl1 expression in mouse endothelium in vivo. (A & B) Analysis of single cell data from Andueza, 2020, PMID 33326796. Single cells collected from low flow (LCA) and normal flow (RCA) intima at either 2 days or 2 weeks post-PCAL induction. (A) Leiden clustering of single cell data. ECs subcluster EC1 – EC7. ECs exposed to normal flow cluster together (EC1–EC3) and those exposed to low flow cluster together (EC4–EC7). Subclusters then separate based on length of exposure, low flow at 2 days (EC4–EC5) and low flow at 2 weeks (EC6–EC7). (B) Violin plots of gene expression of Elavl1, flow- responsive Klf2, and EC inflammatory genes Icam1 and Vcam1. (B) ****p<0.0001, by Wilcoxon rank test.

**SI Figure 5.**
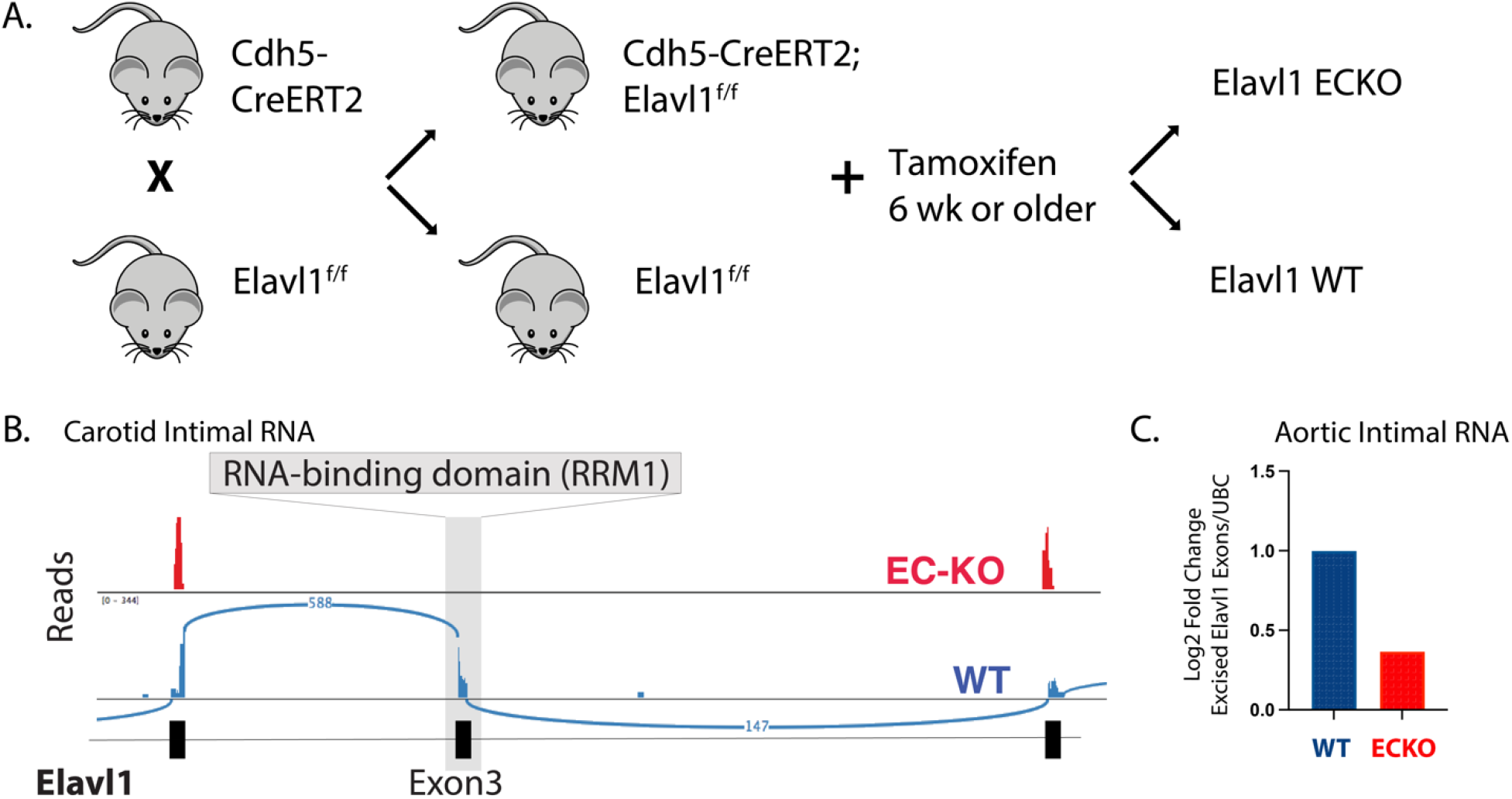
Tamoxifen-induced recombination of Cdh5-CreERT2; Elavl1f/f and Elavl1f/f littermate controls decreased levels of excised region of Elavl1 mRNA. (A) Schematic of generation of experimental mice. (B) Representative RNA reads from an example littermate pair from intimal EC trizol flush. (C) RNA expression levels of excised Elavl1 exons from FACS-isolated mAECs detected by qPCR, normalized to UBC.

**SI Figure 6.**
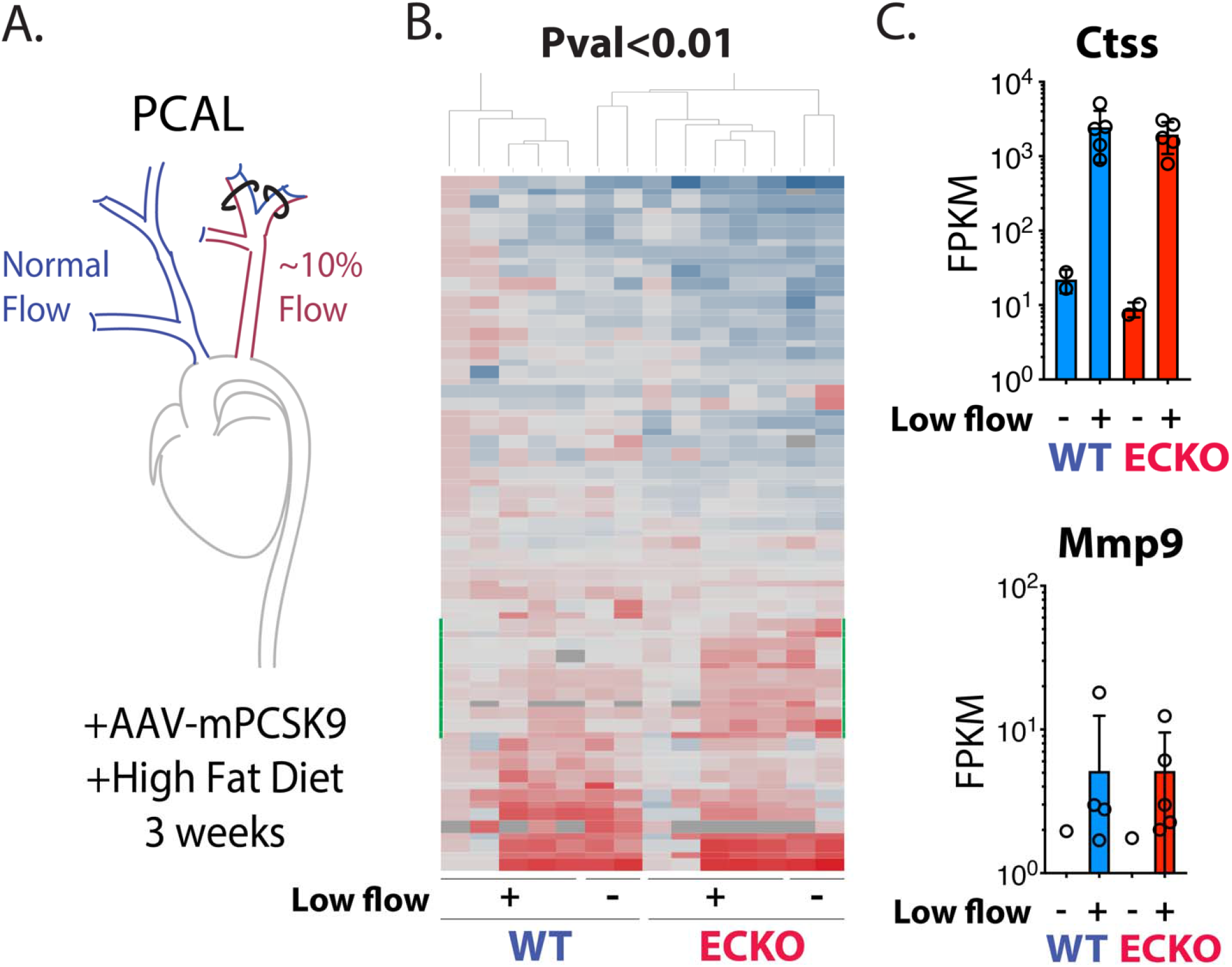
Elavl1 deletion leads few changes in canonically 3’utr stabilized transcripts in bulk endothelial transcriptome. (A) Schematic of PCAL model. Total intimal EC RNA was isolated from low flow LCA and normal flow RCA after three weeks of PCAL and hyperlipidemia via trizol flush. (B&C) Quantification of transcript levels from arterial intima of WT or Elavl1 ECKO mice. Analysis by RSEM and DESeq2 (B) Heat map showing the expression levels of genes for all transcripts achieving P value <0.01. (C) Expression levels of Elavl1 canonically stabilized targets, Ctss and Mmp9. (B&C) N=5 and 5 biological replicates of Elavl1 ECKO and WT mice under low flow, N=2 and 2 under normal flow.

**SI Figure 7.**
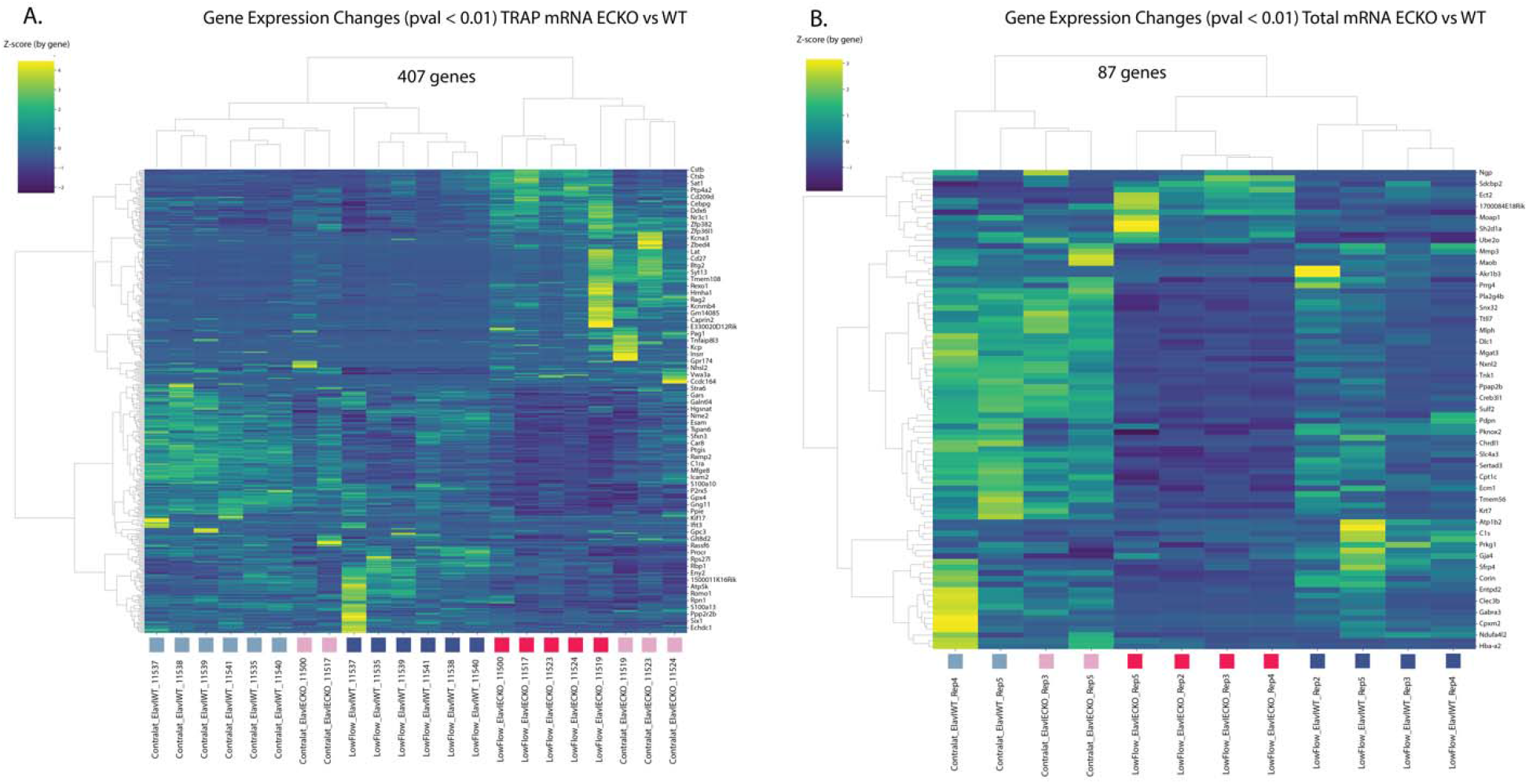
Effects of Elavl1 deletion on TRAP and Total mRNA. Figure shows the transcripts differentially regulated (unadjusted P<0.01) in ribosome-associated RNA (A) and total RNA (B). Mice from which the RNA was isolated are indicated below, with coloring to indicate the genotype (blue = WT and red = ECKO), and shading to indicate the flow pattern (dark = low flow, ligated and light = normal flow, contralateral).

**SI Figure 8.**
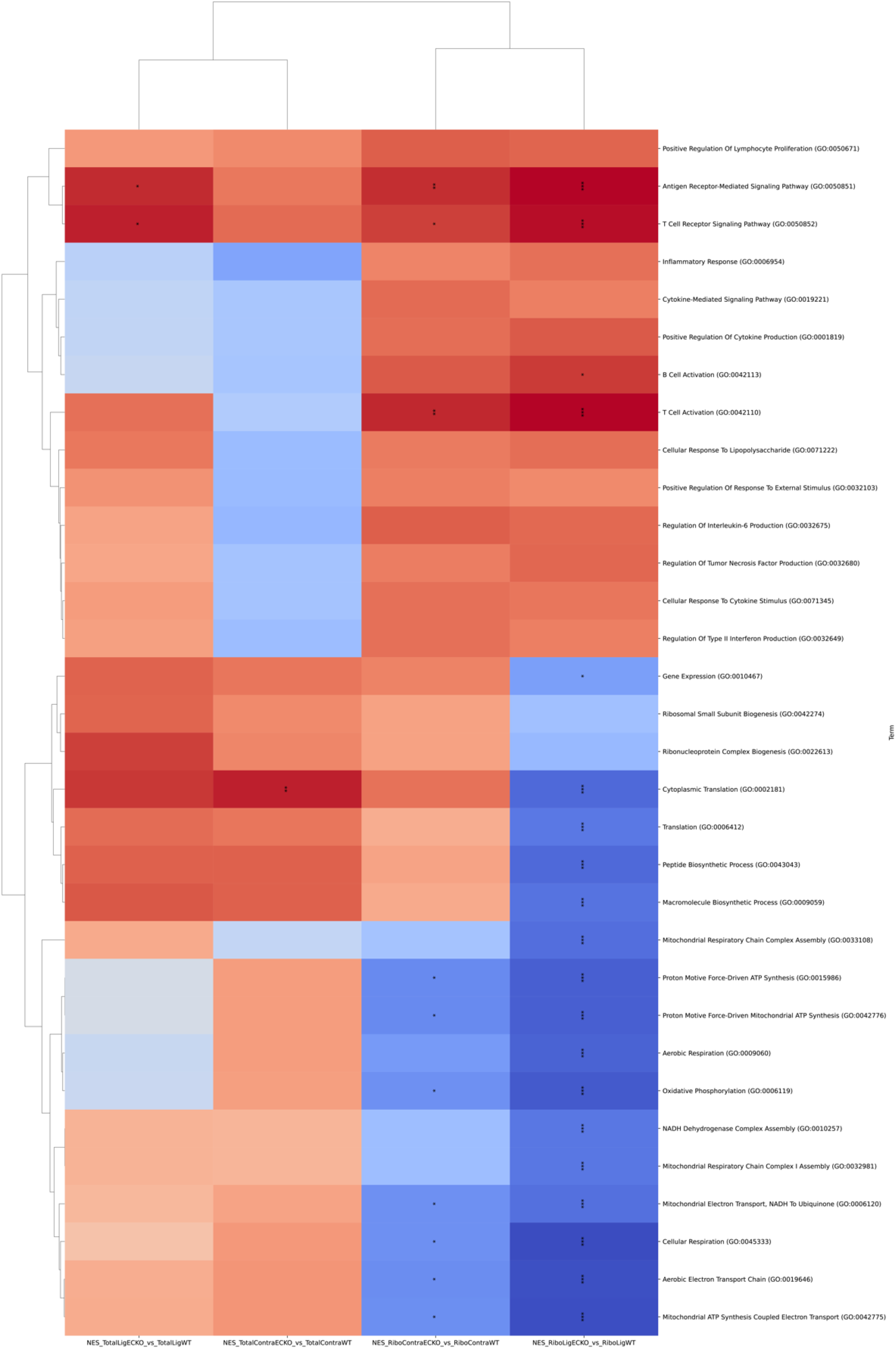
GSEA Terms among transcripts regulated differentially between ECKO and WT mice. Companion figure to main Figure 2J, showing the individual terms different between each response (GSEA weight 1). Plotted colors with the same Normalized Enrichment Score (NES) scale shown in main Figure 2J. *False Discovery Rate (FDR)<0.05, **FDR<0.01, ***FDR<0.001.

**SI Figure 9.**
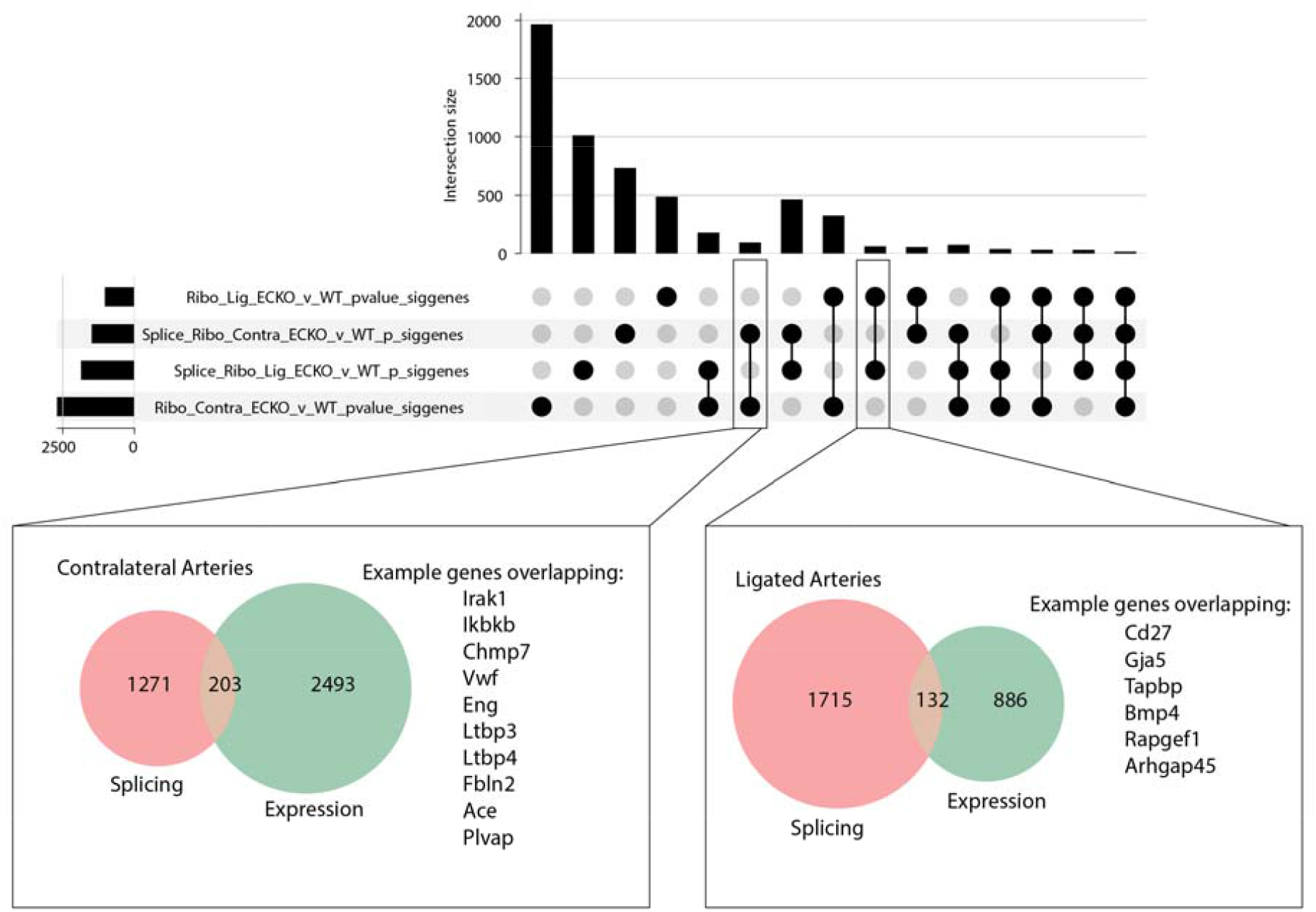
Overlap between genes regulated by splicing or by expression in ECKO mice. Upset plot and Venn diagram of specific examples, showing the overlap between genes regulated by splicing (Leafcutter, p<0.05) or by level at the ribosome (DESeq2, p<0.05). Example genes in overlap are shown adjacent to the Venn diagram.

**SI Figure 10.**
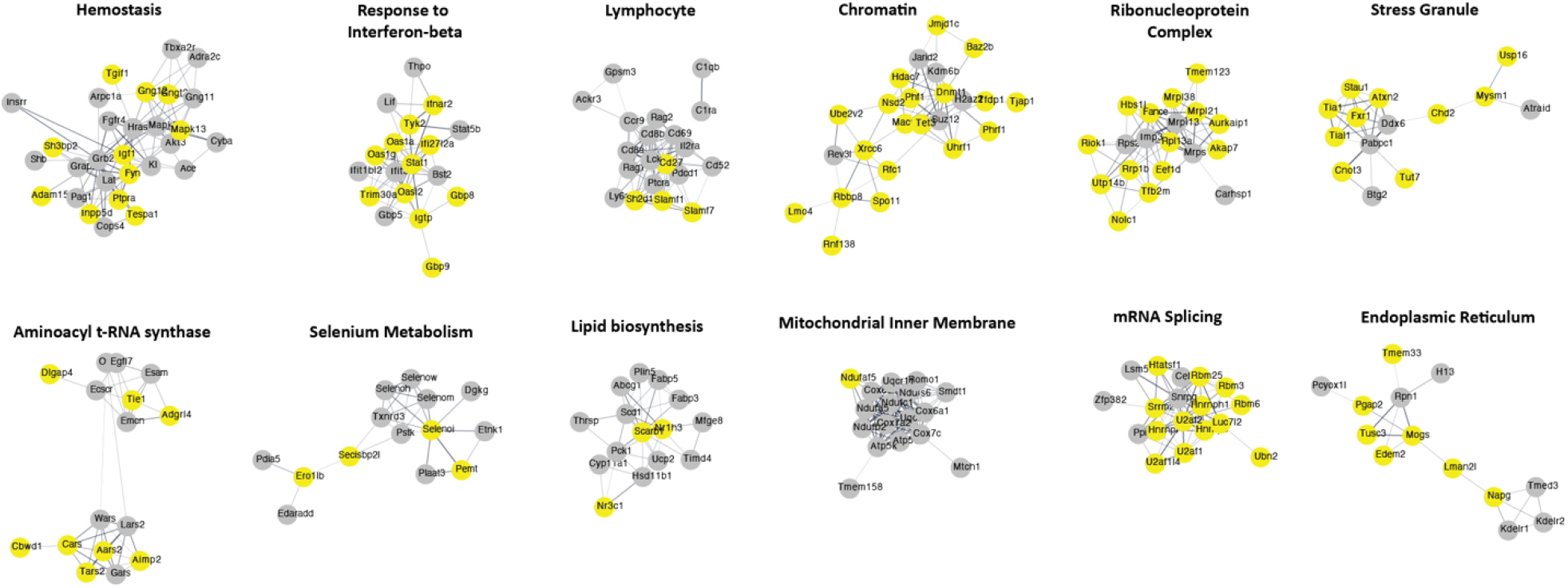
Pathways enriched among TRAP-mRNA changes associated with loss of Elavl1 in low flow endothelium. Cytoscape visualization of all terms associated with low flow transcripts regulated by Elavl1 ECKO in ribosomal mRNA, either by expression (gray; padj <0.05) or splicing (yellow; padj <0.05).

**SI Figure 11.**
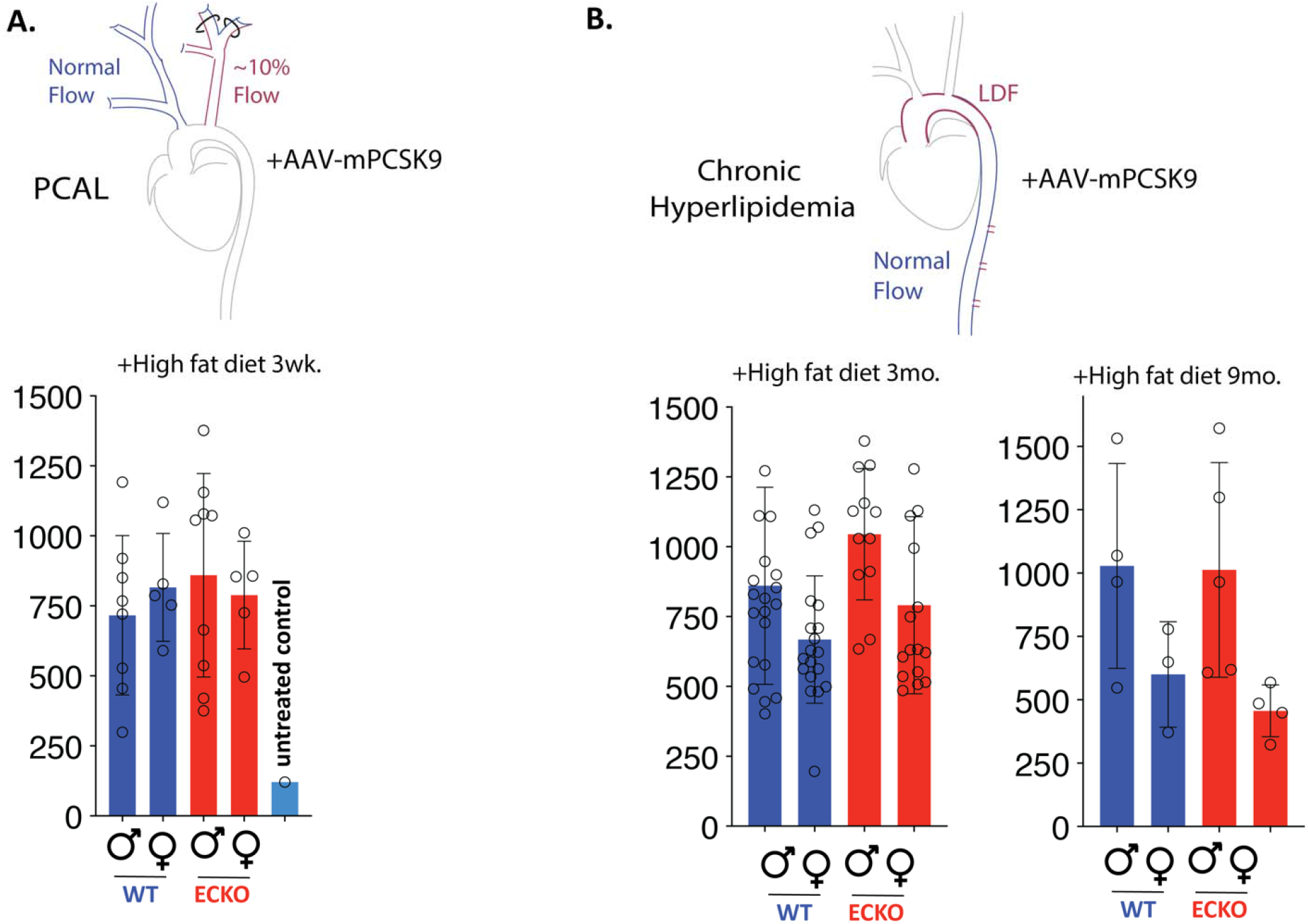
Serum lipid levels are similar in ECKO and WT controls. (A&B) Blood serum total cholesterol levels (mg/dL) at time of collection for PCAL and chronic plaque model, separated by sex and WT vs Elavl1 ECKO.

**SI Figure 12.**
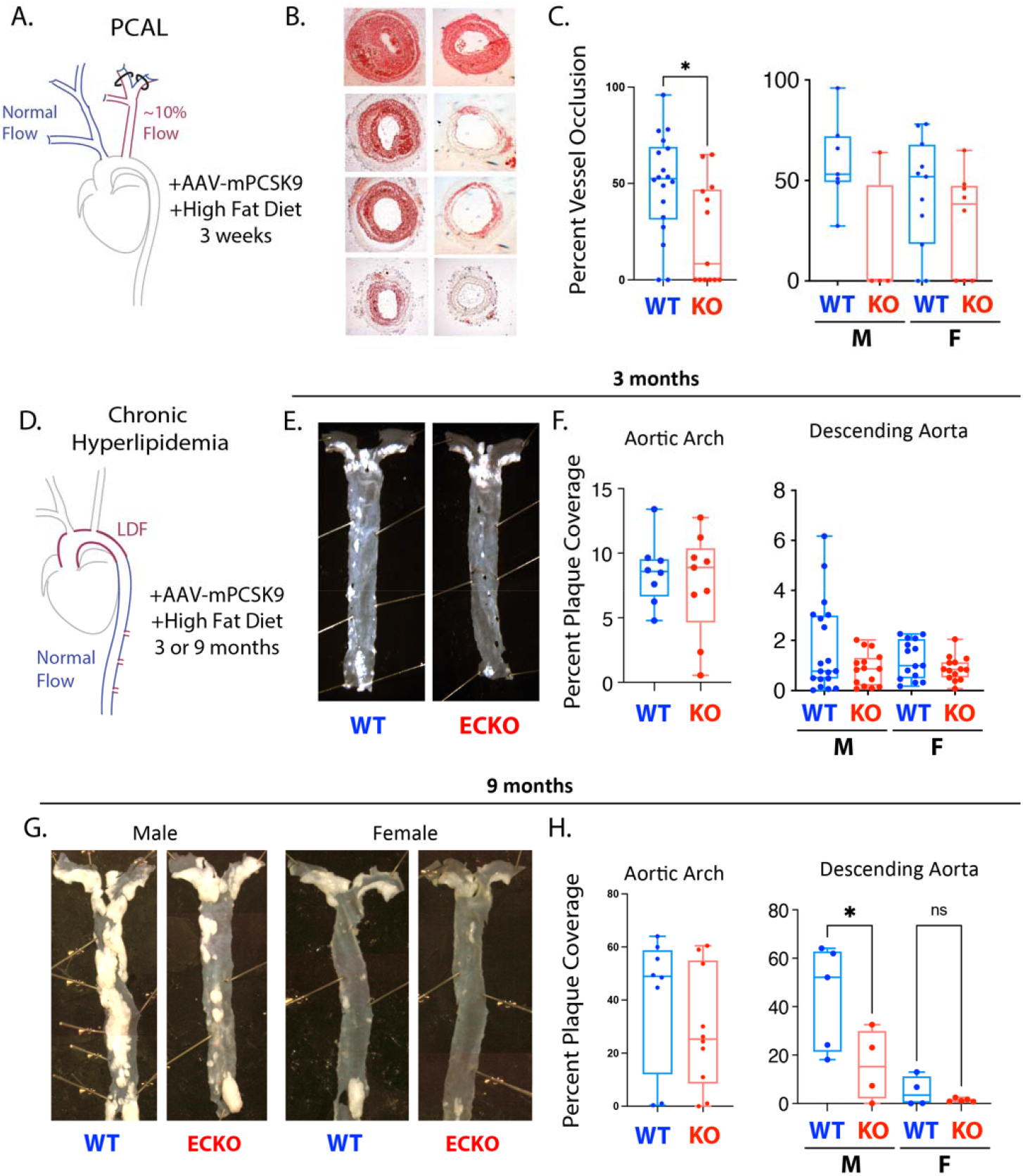
Plaque development in PCAL and at sites of disturbed flow in chronic hyperlipidemia is reduced in Elavl1 ECKO mice. Analysis of plaque coverage by sex in PCAL and chronic plaque development (3 mo and 9 mo hyperlipidemia). (A–C) analysis of percent carotid occlusion at 3 weeks after low flow and hyperlipidemia, with images stained by OilRedO (B) and quantitation (C). (A–H) Similar analysis of en face plaque in chronic response at 3 mo. (D–F) and 9 mo. (G–H).

**SI Figure 13.**
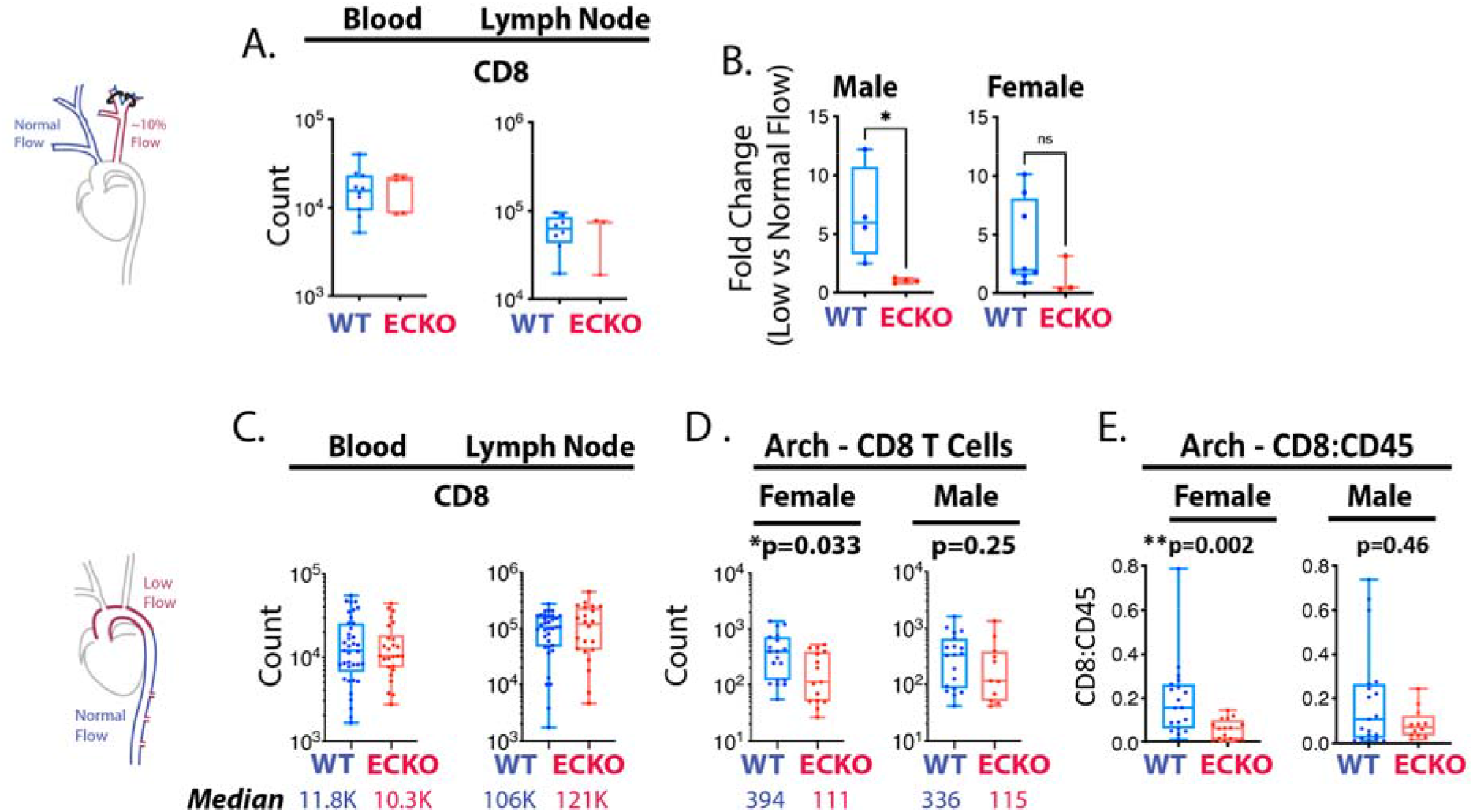
Elavl1 ECKO reduces plaque CD8 T cells without substantial effects on blood or lymphnode levels. (A–E) Quantification of flow cytometry analysis. (A) Absolute count of CD8 T cells in blood and lymph node in mice with PCAL. (B) Ratio of CD8 T cells in low flow atherogenic artery relative to contralateral control artery, separated by sex. (C-D) Absolute count of CD8 T cells with 3 months of hyperlipidemia in blood and lymph node in mice (C) and aortic arch (D). (E) Ratio of CD8:CD45 in aortic arch in female and male mice. (A) N=12 WT, N=7 EC-KO (B) N=8 Female WT, N=4 Male WT; N=3 Female ECKO, N=4 Male ECKO. (C&D) N=37 WT, N=26 ECKO. (E) N=19 Female WT, N=19 Male WT; N=15 Female ECKO, N=11 Male ECKO. Comparisons by two-tailed Mann-Whitney test.

**SI Figure 14.**
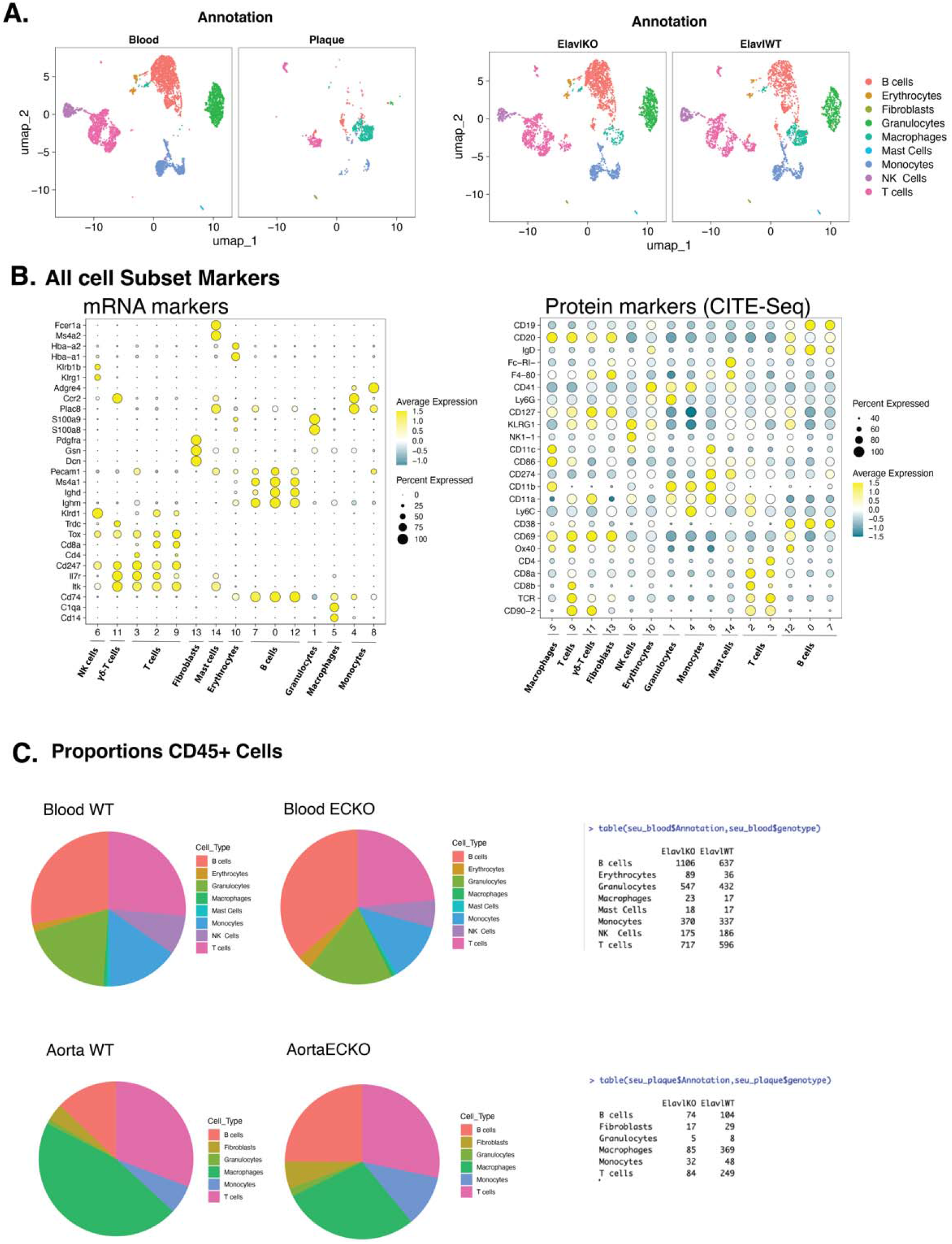
CITE-seq analysis of hematopeotic cells in blood and established aortic plaque of ECKO mice. Graphs showing data analysis of 10x CITE-seq from blood and CD45+ cells from aorta, hashtagged in one 10x reaction. Equal amounts of red-blood cell lysed blood samples were included, and all of the CD45+ aortic cells from each aorta were combined. Ratios between blood samples and between plaque sample therefore reflect in vivo ratios. (A) UMAP separated by tissue source and (B) genotype. (B) Dot plot showing the expression of top RNA and protein markers in hematopoietic subsets. (C) Pie charts showing the percent contribution and total numbers of each cell type in blood and aortic tissue.

**SI Figure 15.**
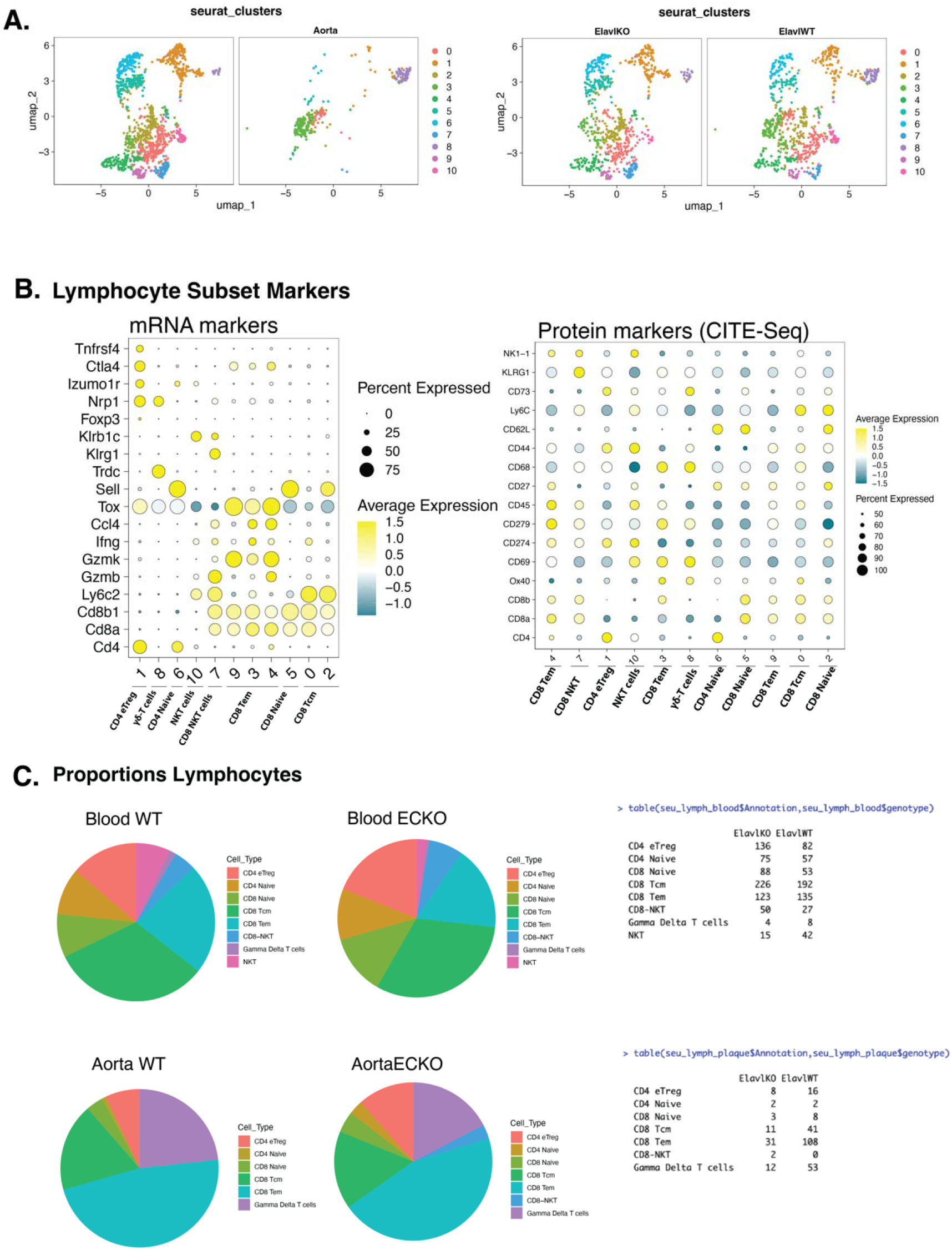
CITE-seq analysis of lymphocytes in blood and established aortic plaque of ECKO mice. Graphs showing data analysis of 10x CITE-seq from blood and CD45+ cells from aorta, hashtagged in one 10x reaction. Equal amounts of red-blood-cell-lysed blood samples were included, and all of the CD45+ aortic cells from each aorta were combined. Ratios between blood samples and between plaque sample therefore reflect in vivo ratios. (A) UMAP separated by tissue source and (B) genotype. (B) Dot plot showing the expression of top RNA and protein markers in lymphocyte subsets. (C) Pie charts showing the percent contribution and total numbers of each cell type in blood and aortic tissue.

**SI Figure 16.**
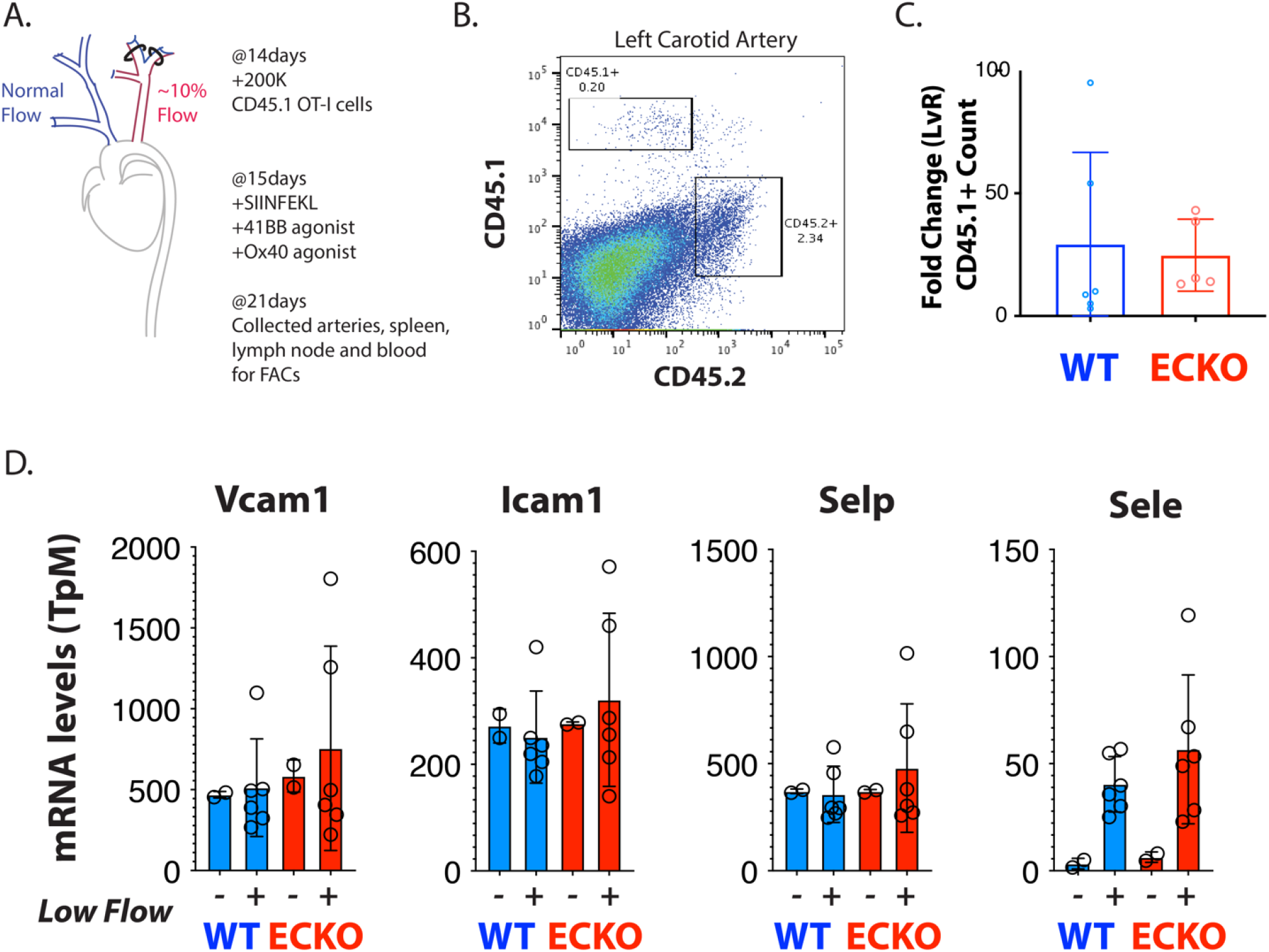
EC Elavl1 does not affect activated CD8+ T cell recruitment to atherosclerotic plaque or expression of adhesion molecules. (A) Schematic of adoptive transfer experimental design. (B) Representative flow cytometry plot featuring detection of both endogenous (CD45.2+) immune cells and adoptively transferred CD45.1+ immune cells in the LCA. (C) Quantification of fold change of CD45.1 numbers recruited to LDF flow LCA over normal flow RCA. (D) mRNA expression (transcript per million) levels of canonical immune cell recruitment molecules in Elavl1 ECKO vs WT in total intimal EC RNAseq data.

**SI Figure 17.**
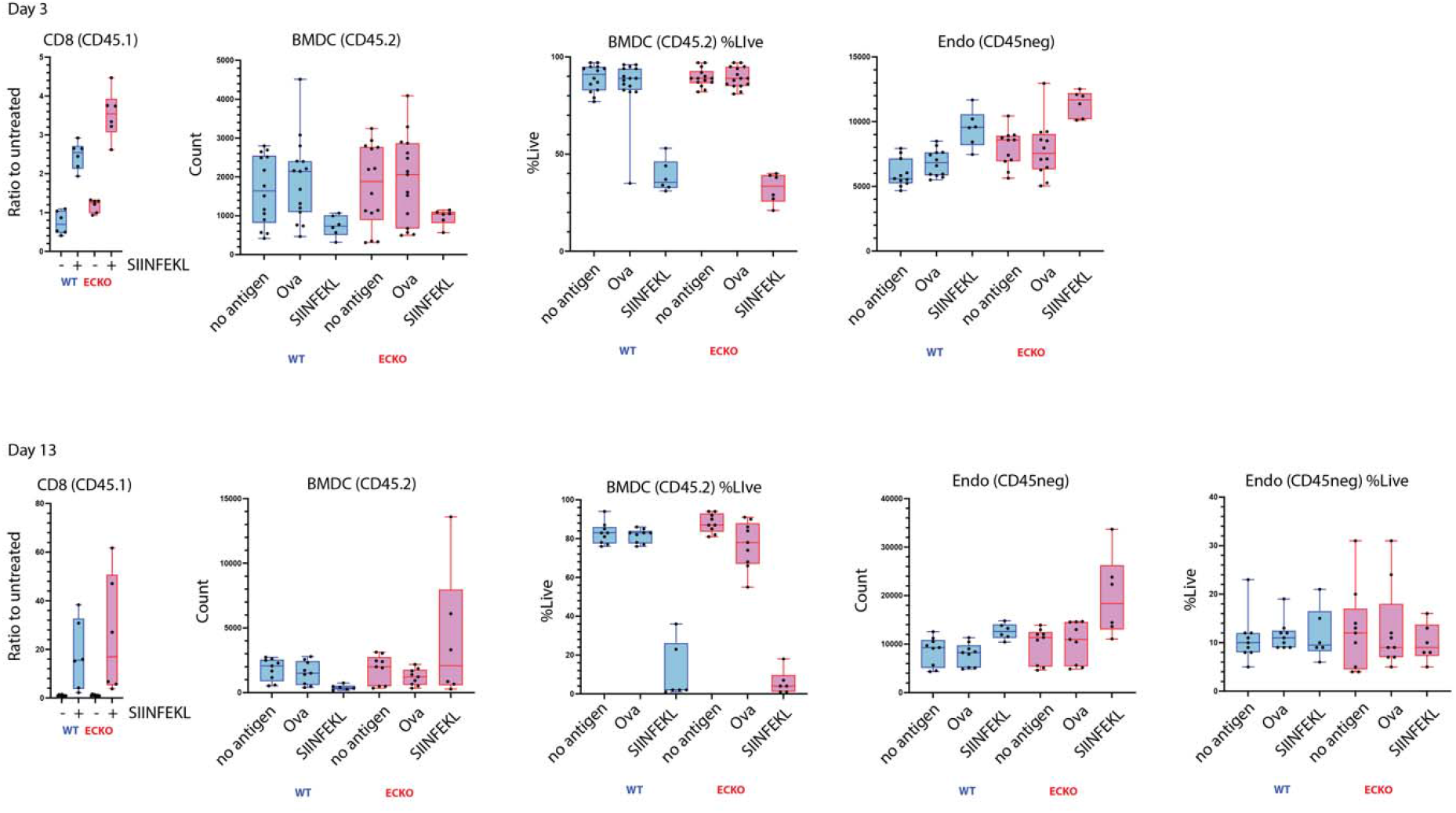
Bone marrow derived dendritic (BMDCs) and endothelial cell numbers are not significantly changed by the loss of Elavl1 in vitro. Graphs show the quantitation of cell numbers of CD45.2+ bone marrow derived dendritic cells, and CD45 negative cells (mainly endothelial, but also including T cells or BMDCs which lost CD45). The BMDCs were either pulsed with SIINFEKL antigen before the addition of T cells (D0), or supplemented with full-length ovalbumin for the indicated period of time (D0 to D3 or D0 to D13). Quantitation is by flow-cytometry, and each point represents cell count from one well of a 96-well plate across multiple individual experimental replicates.

